# DNA demethylation switches the drivers of Foxp3 expression to maintain regulatory T cell identity

**DOI:** 10.1101/2020.12.16.423137

**Authors:** Jun Li, Beisi Xu, Xinying Zong, Minghong He, Yiping Fan, Richard Cross, Jacob H. Hanna, Yongqiang Feng

**Affiliations:** Department of Immunology, St. Jude Children’s Research Hospital, Memphis, TN 38105; Center for Applied Bioinformatics, St. Jude Children’s Research Hospital, Memphis, TN 38105; Department of Molecular Genetics, Weizmann Institute of Science, Rehovot 7610001, Israel

**Keywords:** Regulatory T cells, DNA demethylation, cell fate, Foxp3

## Abstract

Maintenance of differentiated cellular states is crucial for numerous biological processes, yet its molecular basis remains unclear. Here, we investigate how mechanistically regulatory T (T_reg_) cell fate is “locked in” during lineage commitment via transcriptional regulation of its lineage-specifying factor Foxp3. Tet-mediated DNA demethylation of *Foxp3* enhancer CNS2 was proposed to be a key mechanism maintaining Foxp3 transcription. However, this model has not been directly tested. Therefore, we integrated genetic, pharmacological, and epigenetic approaches to examine the function and mechanism of DNA demethylation in T_reg_ lineage maintenance. We observed an abrupt switch of the transcriptional drivers of Foxp3 upon DNA demethylation, which was abolished by CNS2 deficiency. Demethylation of CNS2 increased chromatin accessibility and protein binding, conferring on T_reg_ fate substantial resistance to adverse environments. Thus, our study consolidated the role of DNA demethylation in stabilizing Foxp3 expression in *cis* and revealed a novel regulatory mode governing Tr_eg_ identity.

## INTRODUCTION

T_reg_ cells are induced from precursor cells by cognate antigens in the presence of favorable environmental cues, including IL-2 and TGF-β, to suppress specific pathogenic T cells and maintain immune homeostasis (Josefowicz, Lu, & Rudensky, 2012; Savage, Klawon, & Miller, 2020). Successful T_reg_ lineage commitment further relies on subsequent mechanisms to “lock in” the T_reg_ fate (Fu et al., 2012; Yue, Lio, Samaniego-Castruita, Li, & Rao, 2019; Yue et al., 2016). Genetic fate mapping and acute depletion experiments indicate that T_reg_ cells are long-lived and that their continuous activity is required to maintain immune tolerance (J. M. Kim, Rasmussen, & Rudensky, 2007; Rubtsov et al., 2010). Thus, a stable lineage in the absence of guidance cues or in the presence of adverse environments is critical for sustained T_reg_ immune suppressive function.

T_reg_ lineage commitment and function relies on nuclear factor Foxp3 (Brunkow et al., 2001; Fontenot, Gavin, & Rudensky, 2003; M. Gavin & Rudensky, 2003; Hori, Nomura, & Sakaguchi, 2003; Liston et al., 2007). Investigation of the mechanisms controlling Foxp3 expression has produced enormous insights into T_reg_ biology (Josefowicz et al., 2012; Sakaguchi et al., 2020; Savage et al., 2020). Although a growing list of cell-extrinsic and -intrinsic factors was known to maintain T_reg_ fate, an epigenetic memory conferred by DNA demethylation was proposed as a key mechanism stabilizing Foxp3 transcription (Polansky et al., 2008). This process is initiated by Tet dioxygenases and accomplished by a multi-step enzymatic reaction. In the absence of *Tet2* and *Tet3*, T_reg_ cells fail to sustain Foxp3 expression (Nakatsukasa et al., 2019; Yue et al., 2019; Yue et al., 2016). Because *Foxp3* enhancer CNS2 is demethylated by Tet enzymes in T_reg_ cells and not in precursors or conventional T cells (Polansky et al., 2008; Yue et al., 2016) and CNS2-deficiency leads to unstable Foxp3 expression (Feng et al., 2014; X. Li, Liang, LeBlanc, Benner, & Zheng, 2014; Zheng et al., 2010), Tet-dependent DNA demethylation was proposed to stabilize Foxp3 transcription through CNS2.

However, Tet enzymes also control the expression of many other genes (Nakatsukasa et al., 2019; Ohkura et al., 2012; Yue et al., 2019), which may stabilize Foxp3 expression in *trans*. It is unclear to what extent Tet-dependent *cis* and *trans* mechanisms contribute to the stability of Foxp3 transcription. A direct test of the sufficiency of CNS2 in mediating Tet-dependent stabilization of Foxp3 expression is lacking. Although several factors have been shown to maintain Foxp3 expression, including IL-2/Stat5 signaling, Runx-CBFβ, Foxp1, Foxo1/Foxo3, and Akt/Pten (Feng et al., 2014; Ghosh, Roy-Chowdhuri, Kang, Im, & Rudra, 2018; Ouyang et al., 2010; Rudra et al., 2009; Shrestha et al., 2015), their relationships with Tet/DNA demethylation have not been fully explored. Addressing these issues will produce important insights into the mechanisms governing T_reg_ lineage stability critical in various immunological settings.

Here, we modeled T_reg_ lineage commitment and maintenance *ex vivo* with mouse primary T cells by taking advantage of vitamin C (or ascorbic acid, ASC)–dependent Tet enzymatic activity (Sasidharan Nair, Song, & Oh, 2016; Yue et al., 2016). We integrated pharmacological, genetic, and epigenetic approaches to assess the role and underlying mechanisms of Tet-dependent DNA demethylation in maintaining qualitative Foxp3 expression in *cis* through enhancer CNS2. Our study uncovered a novel regulatory mode established by hypomethylated CNS2, which switches the transcriptional drivers of Foxp3 expression to maintain T_reg_ identity.

## RESULTS

### Histone acetylation drives both Foxp3 induction efficiency and expression levels during early T_reg_ cell development

We first sought to uncover the driving force of Foxp3 transcription during T_reg_ cell development. To this end, we reexamined permissive histone acetylation (e.g., H3K27ac) in Foxp3 induction because of its essential role in promoting gene expression (Fujisawa & Filippakopoulos, 2017). H3K27ac is deposited by histone acetyltransferases (HATs) and read by bromodomain (BRD)-containing proteins (Fujisawa & Filippakopoulos, 2017). Using Cut&Run sequencing (Fig. 1*A*), which probes epigenetic modifications in their native states with improved resolution (Skene & Henikoff, 2017), we confirmed a broad H3K27ac modification around the *Foxp3* locus in *ex vivo*–isolated T_reg_ (nT_reg_) cells but not in CD4 naïve T (Tn) cells. We then used pharmacological inhibitors to examine the stage-specific roles of histone acetylation during T_reg_ cell differentiation (iT_reg_) from CD4 Tn cells *in vitro*, which recapitulates T_reg_ cell development *in vivo* (Feng et al., 2015; Josefowicz et al., 2012; Rudensky, 2011; Yue et al., 2019). Blockade of HAT p300/CBP with C646, I-CBP112, or SGC-CBP30 reduced Foxp3 induction efficiency in a dose-dependent manner (Figs. 1*B* and S1*A, B*). To test whether BRD proteins are required to transduce the histone acetylation signal, we treated differentiating T cells with BRD inhibitors JQ1, I-BET151, bromosporine, or PFI-1, resulting in a dose-dependent inhibition of Foxp3 induction efficiency (Figs. 1*C* and S1*C*-*E*). These results confirmed the essential role of histone acetylation signaling in driving Foxp3 induction during T_reg_ cell development (Feng et al., 2015; Liu et al., 2014).

**Fig. 1.**
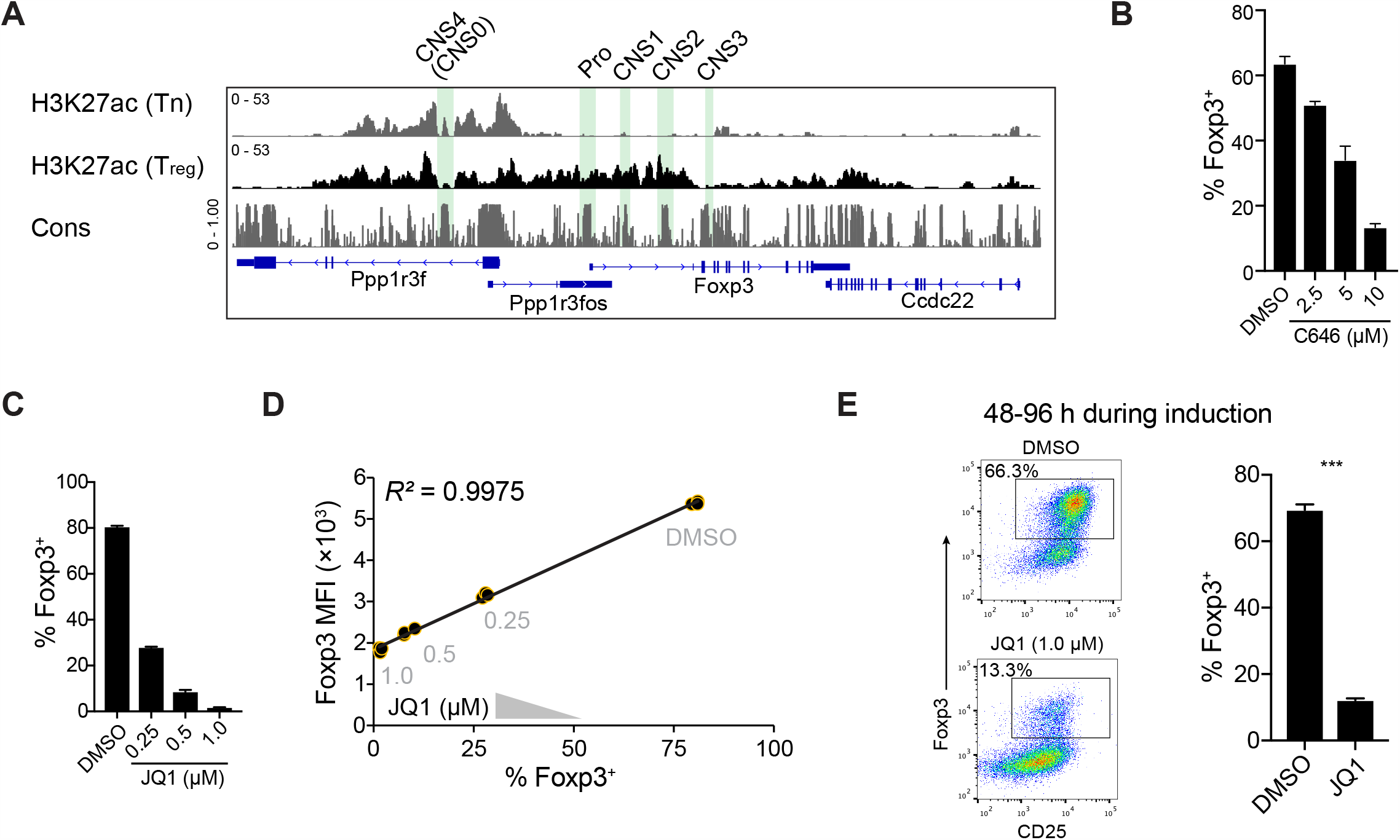
Histone acetylation drives both Foxp3 induction efficiency and expression levels during early T_reg_ cell development. *(A)* Quantification of histone H3K27ac modification around the *Foxp3* locus (mm9) in CD4 Tn and nT_reg_ cells by Cut&Run-seq. Positions of the *Foxp3* promoter (Pro) and enhancers are highlighted. Cons, DNA sequence conservation among placental mammals. *(B)* Foxp3 induction efficiency in the presence of titrated C646 during T_reg_ cell development *in vitro*. CD4 Tn cells were cultured in T_reg_-induction conditions (TCR agonists, IL-2, and TGF-β) with or without C646 for 4 days before analyzing Foxp3 expression. Data depict means + SEMs of triplicates and represent 3 independent experiments. (*C, D*) Foxp3 induction efficiency in the presence of titrated JQ1 during T_reg_ cell development *in vitro* (*C*). Correlation of Foxp3 induction efficiency and Foxp3 expression levels per cell (median fluorescence intensity, MFI) was calculated (*D*). Data depict means + SEMs of triplicates and represent 3 experiments. (*E*) Inhibition of BRD proteins silenced Foxp3 expression at the late stages of T_reg_ cell development. JQ1 (1 µM) was added after CD4 Tn cells were cultured in T_reg_-induction media for 2 days. Foxp3 expression was analyzed on day 4. Data depict means + SEMs of triplicates and represent 2 independent experiments. \*\*\**p*<0.001 (unpaired, two-tailed *t* test). See also Fig. S1.

Because T_reg_ lineage determination is a qualitative trait and Foxp3 expression levels are a quantitative one, we reasoned that these two categories of gene regulation could be controlled by separate mechanisms. Therefore, we tested whether histone acetylation signaling plays different roles in these two potentially distinct regulatory modes. In the presence of titrated amounts of HAT or BRD inhibitors (e.g., JQ1), the relationship between Foxp3 induction efficiency and Foxp3 expression levels demonstrated that qualitative and quantitative controls of Foxp3 expression were well correlated (*R*^*2*^=0.9975; Fig. 1*D*), suggesting that, at the early stage of T_reg_ cell development, Foxp3 induction efficiency and Foxp3 expression levels could be controlled by the same mechanisms. Because blockade of HATs or BRD proteins suppresses global gene expression during T cell activation and differentiation, thereby inhibiting Foxp3 expression, we then added JQ1 into culture media 2 days after CD4 Tn cells being cultured in T_reg_-induction conditions. This acute blockade of BRD proteins abruptly reduced the number of Foxp3-expressing cells (Fig. 1*E*), and the remaining Foxp3^+^ cells showed significantly lower levels of Foxp3 and its target genes (e.g., CD25, GITR, and CTLA-4) (Fig. S1*F*). These results indicate that histone acetylation signaling determines both Foxp3 induction efficiency and expression levels during early T_reg_ cell development.

### Tet enzymes switch the driving force of qualitative Foxp3 expression

After stochastic induction, Foxp3 expression is maintained in differentiated T_reg_ cells during their long lifespan. Tet-dependent DNA demethylation was proposed to be the main mechanism acting in *cis* to “lock in” Foxp3 expression in committed T_reg_ cells. Yet, the nature of the transition from Foxp3 induction to maintenance remains unclear. Because this process is not readily accessible in T_reg_ cell differentiating *in vivo* (Fig. 2*A*), we modeled T_reg_ lineage commitment *in vitro* using mouse CD4 Tn cells in the presence or absence of Tet-induced DNA demethylation (Sasidharan Nair et al., 2016; Yue et al., 2016).

**Fig. 2.**
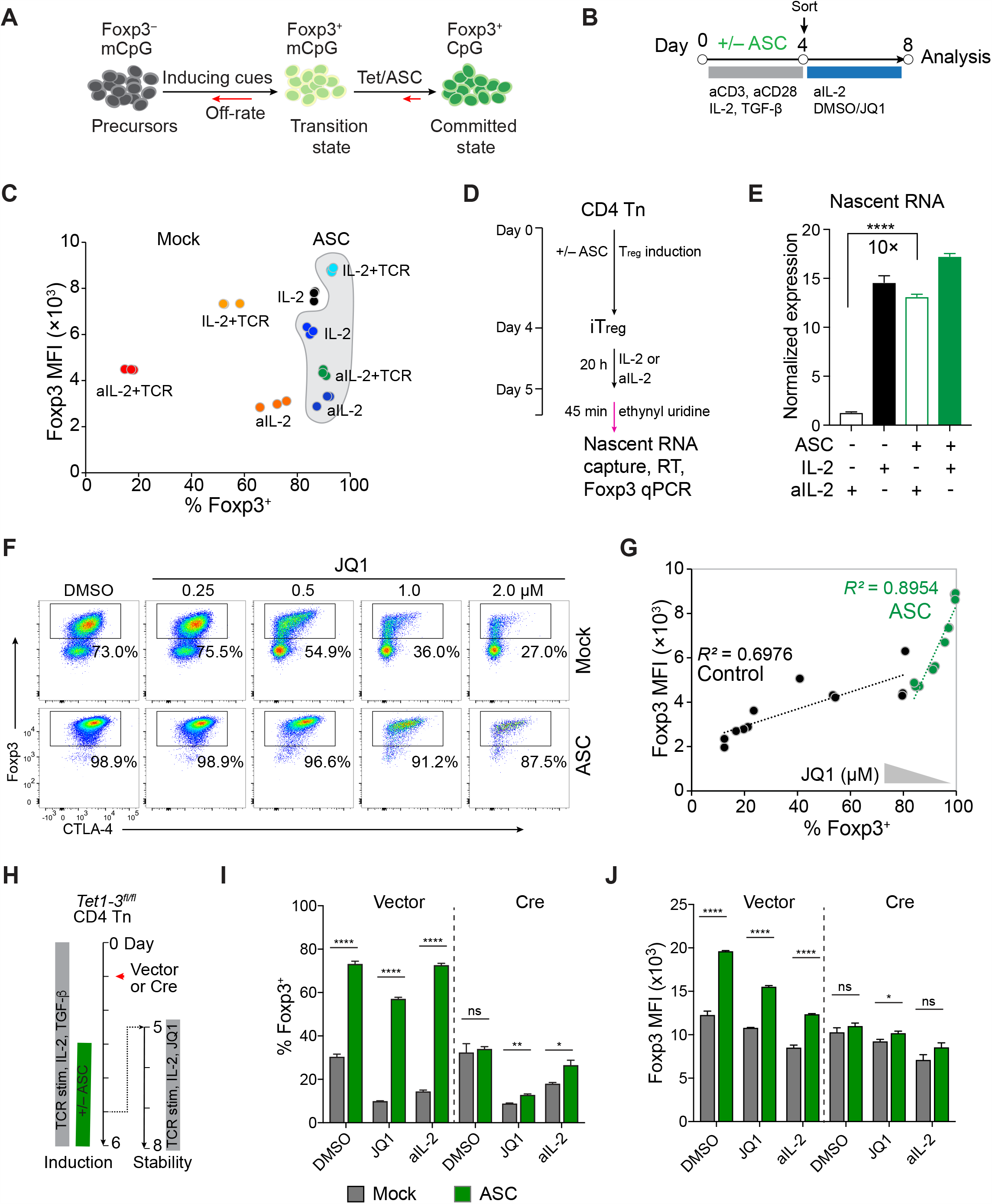
Tet enzymes switch the driving force of qualitative Foxp3 expression. *(A)* Schematic processes of T_reg_ lineage commitment. Foxp3 expression is triggered by T_reg_-induction cues and stabilized by ASC-dependent Tet enzymatic activity. mCpG, methylated CpG. The off rate indicates the instability of Foxp3 expression. *(B)* Experimental procedures for assaying the stability of Foxp3 expression *in vitro*. On day 4, iT_reg_ cells induced with or without supplemented ASC were isolated and then grown in media containing recombinant IL-2, IL-2 neutralization antibodies (aIL-2), or JQ1 with or without TCR restimulation for an additional 4 days before analysis. *(C)* Stability and levels of Foxp3 expression in mock- and ASC-pretreated (gray shaded) iT_reg_ cells grown in media containing recombinant IL-2 or aIL-2 with or without TCR restimulation by plate-bound anti-CD3 and anti-CD28 antibodies. Dots indicate technical replicates. Data represent 2 experiments. (*D, E*) Quantification of transient Foxp3 transcription. Mock- and ASC-treated iT_reg_ cells were cultured with IL-2 or aIL-2 for 20 h and then pulse-chased with ethynyl uridine (EU) for 45 min before quantification of EU-labeled nascent Foxp3 mRNA via RT-qPCR. Signals of EU-enriched mRNA were normalized to those of input cells in each sample. \*\*\*\**p*<0.0001 (two-tailed, unpaired *t* test). (*F, G*) Stability and levels of Foxp3 expression in mock- and ASC-pretreated iT_reg_ cells grown in media containing titrated JQ1 for an additional 4 days. Data represent more than 3 independent experiments. (*H*-*J*) *Tet1-3*^*fl/fl*^ CD4 Tn cells were cultured in T_reg_-induction medium before being transduced with Cre-expressing retrovirus on day 1. PBS or ASC was added from day 3 to day 5. Cells were then grown in media containing DMSO (with IL-2), aIL-2, or JQ1 (with IL-2) with TCR restimulation for an additional 3 days before analysis. Data show means + SEMs of triplicates and represent one of 2 independent experiments. ns, not significant; \**p*<0.05, \*\**p*<0.01, \*\*\*\**p*<0.0001 (two-tailed, unpaired *t* test). See also Fig. S2.

We isolated CD4 Tn cells from wild-type (WT) *Foxp3*^*gfp*^ knock-in mice and induced iT_reg_ cells in media supplemented with or without ASC. To assess the stability or qualitative regulation of Foxp3 expression, we sorted iT_reg_ cells by GFP expression 4 days later and cultured them in media containing recombinant IL-2, IL-2 neutralization antibodies (aIL-2), or JQ1, with or without TCR restimulation by anti-CD3 and anti-CD28 antibodies (Fig. 2*B*). More than 80% of mock-treated iT_reg_ cells lost Foxp3 expression after TCR restimulation and IL-2 deprivation, whereas more than 50% of cells maintained Foxp3 expression in the presence of IL-2 or absence of TCR restimulation (Fig. 2*C*). Similar to stable nT_reg_ cells, more than 80% of ASC-treated iT_reg_ cells remained Foxp3^+^ regardless of the culture conditions (Figs. 2*C* and S2*A*). Notably, Foxp3 expression levels were still controlled by IL-2 and TCR signaling in ASC-treated iT_reg_ cells. These results strongly suggest that, after ASC treatment, qualitative and quantitative Foxp3 expression are controlled by distinct mechanisms. To understand the features of Foxp3 transcription during this transition, we labeled nascent RNA with pulsed ethynyl uridine to directly compare the transient transcriptional activity of Foxp3 in mock-treated iT_reg_ cells to that in ASC-treated iT_reg_ cells. We found that Foxp3 transcription was immediately silenced upon IL-2 deprivation in mock-treated iT_reg_ cells but not in ASC-treated iT_reg_ cells (Fig. 2*D, E*). Together, these results indicate that ASC treatment induced a drastic change of the mechanism governing qualitative Foxp3 transcription (e.g., from IL-2–sensitive to IL-2–insensitive regulations).

Next, we examined the role of histone acetylation signaling in stable Foxp3 expression. We sorted mock- and ASC-treated iT_reg_ cells and cultured them in media containing titrated amounts of JQ1, which abruptly silenced Foxp3 expression in mock-treated iT_reg_ cells (Fig. 2*F*). In contrast, Foxp3 expression was maintained in more than 85% of ASC-pretreated iT_reg_ cells. Akin to IL-2 deprivation, JQ1 reduced Foxp3 median fluorescence intensity (MFI) in both mock- and ASC-pretreated iT_reg_ cells (Fig. 2*G*). These data further indicate a profound transcriptional reprogramming of the qualitative regulation of Foxp3 expression after ASC treatment.

To test whether Tet enzymes mediate the effect of ASC, we acutely ablated *Tet1, Tet2*, and*Tet3* (*Tet1-3*) in CD4 Tn cells isolated from *Tet1-3*^fl/fl^ mice with Cre-expressing retrovirus (Fig. 2*H*). Because of the short time window of cell culture, our approach did not completely deplete Tet proteins (Fig. S2*B*). Nonetheless, Cre-transduced iT_reg_ cells significantly decreased the stability of Foxp3 expression in ASC-pretreated iT_reg_ cells after being exposed to JQ1 or upon IL-2 neutralization (Fig. 2*I*). Ablation of Tet proteins also reduced ASC-dependent upregulation of Foxp3 expression (Fig. 2*J*). Neither ASC treatment nor Tet deficiency influenced Foxp3 induction efficiency (Fig. S2*C*). Histone acetylation signaling and Tet activity coordinated to upregulate Foxp3 and CD25 expression levels in iT_reg_ cells (Figs. 2*J* and S2*D*). The latter may contribute to Tet/ASC–dependent stabilization of Foxp3 expression in *tran*s. Our results are consistent with the reported function of ASC in maintaining T_reg_ identity through Tet enzymes (Sasidharan Nair et al., 2016; Yue et al., 2016). Importantly, we observed that Tet enzymes drastically switched the drivers of qualitative Foxp3 transcription, conferring stable T_reg_ identity in adverse environments.

### Tet enzymes stabilize qualitative Foxp3 expression via enhancer CNS2

Next, we examined the requirement for CNS2 in ASC-dependent stabilization of Foxp3 expression (Fig. 3*A*). We first performed whole-genome bisulfite sequencing (WGBS) to fine-map the differential DNA methylation patterns in CD4 Tn, effector T (Te), nT_reg_, and iT_reg_ cells developed with or without supplemented ASC. Because the *Foxp3* gene is on the X chromosome that undergoes random inactivation, cells used for WGBS were isolated from male *Foxp3*^*gfp*^ mice. Although global DNA methylation ratios were not drastically different (Fig. S3*A*), differentially methylated regions (DMRs) unambiguously distinguished these cell types, as illustrated by the principal component analysis (Fig. S3*B*). Compared to CD4 Tn and Te cells, nT_reg_ cells were hypomethylated at the *Foxp3* promoter, CNS2, and several other regions in the locus (Fig. 3*B*). ASC treatment induced hypomethylation of these DMRs in iT_reg_ cells to a level comparable to that in nT_reg_ cells despite reduced widths. Notably, many T_reg_-specific DMRs, such as *Lrrc32, Emsy, Il2ra, Ift80, Wdr82*, and *Ctla4*, were less hypomethylated in ASC-treated iT_reg_ cells (Fig. S4*A*). Although the cause of these differences was unknown, our results (Figs. 3*B* and S4*B-G*) together with published reports (Sasidharan Nair et al., 2016; Yue et al., 2016) indicate that ASC-treated iT_reg_ cells serve as an excellent *ex vivo* model in which to study the mechanistic processes of DNA demethylation–dependent control of Foxp3 transcription during T_reg_ lineage commitment.

**Fig. 3.**
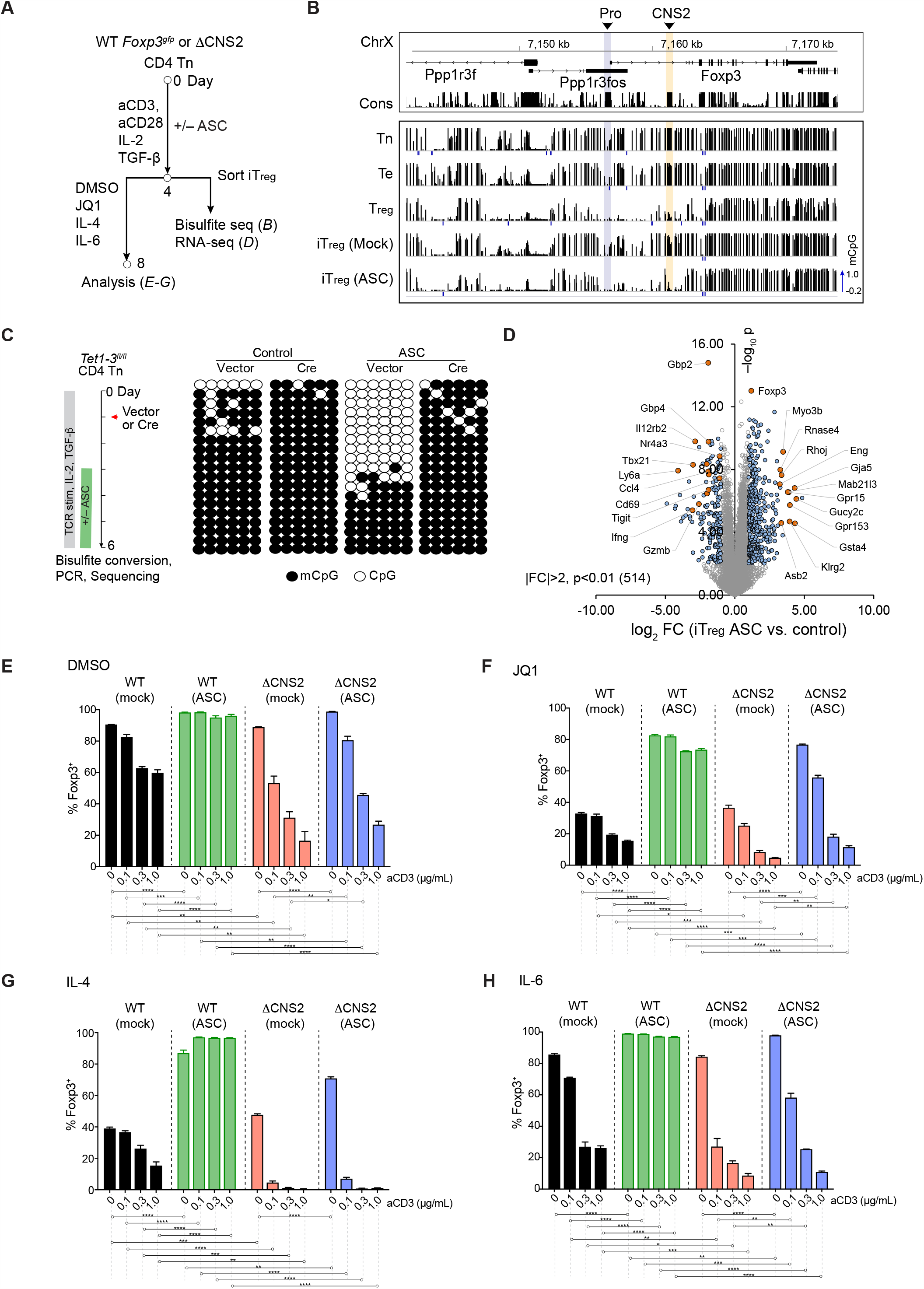
Tet enzymes stabilize qualitative Foxp3 expression via enhancer CNS2. *(A)* Experimental procedures of assaying T_reg_ stability, differential gene expression, and WGBS with iT_reg_ cells induced from *Foxp3*^*gfp*^ CD4 Tn cells with or without supplemented ASC. *(B)* DNA (CpG) methylation ratios around the *Foxp3* locus (mm9) in CD4 Tn cells, effector T (Te) cells, nT_reg_ cells, and mock- or ASC-treated iT_reg_ cells. Vertical lines represent individual CpG sites and methylation levels from 0 (unmethylated) to 1 (methylated). Data were merged from 2 biological replicates. Regions covered by < 5 reads are marked as −0.2. *(C)* Quantification of the methylation status at select CpG sites of CNS2 by bisulfite PCR-seq in iT_reg_ cells after acute deletion of Tet enzymes followed by mock and ASC treatment. Rows of dots represent individual DNA molecules. *(D)* Genes that are differentially expressed in mock- and ASC-treated iT_reg_ cells. Data were merged from 2 biological replicates. (*E*-*H*) Stability of Foxp3 expression in WT *Foxp3*^*gfp*^ and *Foxp3*^*ΔCNS2-gfp*^ iT_reg_ cells pretreated with or without ASC. Cells were then restimulated with titrated amounts of plate-bound anti-CD3 antibody and constant anti-CD28 antibody (1 µg/mL) in media containing recombinant IL-2 (100 U/mL) and either DMSO (*E*), JQ1 (2 µM) (*F*), IL-4 (10 ng/mL) (*G*), or IL-6 (20 ng/mL) (*H*) for 4 days before analysis. \**p*<0.05, \*\**p*<0.01, \*\*\**p*<0.001, \*\*\*\**p*<0.0001 (unpaired, two-tailed *t* test). Data show means + SEMs of triplicates and represent 3 experiments. See also Figs. S3 and S4.

We then tested whether ASC-induced hypomethylation of CNS2 was indeed Tet-dependent. To this end, we acutely ablated *Tet1-3* with retroviral Cre during T_reg_ cell development, which abolished the effect of ASC on CNS2 demethylation (Fig. 3*C*), consistent with the reported role of Tet/ASC in CNS2 demethylation (Sasidharan Nair et al., 2016; Yue et al., 2016). To uncover the global effect of ASC treatment, we performed RNA-seq with mock-and ASC-treated iT_reg_ cells, revealing 514 genes that were up- or down-regulated (*p*<0.01 and fold change [FC] >2; Fig. 3*D*). Although these genes’ exact functions remain to be determined, this result suggests potential *trans* mechanisms controlling the stability of Foxp3 expression.

To test this possibility, we assessed whether CNS2 was indeed responsible for Tet/ASC-dependent stabilization of Foxp3 expression. We induced iT_reg_ cells from CNS2-sufficient and -deficient CD4 Tn cells in media supplemented with or without ASC. We then sorted iT_reg_ cells after induction and examined the stability of Foxp3 expression in adverse environments, including IL-4, IL-6, or JQ1, in the presence of titrated amounts of TCR agonists. In mock-treated iT_reg_ cells, TCR restimulation alone or in combination with JQ1, IL-4, or IL-6 drastically diminished the percentages of Foxp3-expressing cells in both CNS2-sufficient and -deficient cells (Fig. 3*E*-H). Notably, mock-treated CNS2-deficient iT_reg_ cells appeared to be less stable in Foxp3 expression than CNS2-sufficient iT_reg_ cells, suggesting that CNS2 may also play a role in maintaining Foxp3 expression before DNA demethylation. In contrast, ASC treatment drastically stabilized Foxp3 expression in WT iT_reg_ cells regardless of the adverse environments. The same treatment failed to stabilize Foxp3 expression in CNS2-deficient iT_reg_ cells, indicating that CNS2 is required for ASC-dependent stabilization of qualitative Foxp3 expression. As CNS2’s main function was enabled by ASC-treatment that relied on Tet enzymes, our results demonstrated the sufficiency of Tet-induced DNA demethylation of CNS2 in stabilizing Foxp3 expression and excluded the significance of potential Tet/ASC-mediated *trans* mechanisms in stabilizing Foxp3 expression. Our experiments also uncovered a novel regulatory mode established by hypomethylated CNS2, predominantly switching the transcriptional driving force of qualitative Foxp3 expression to stabilize T_reg_ lineage identity.

### DNA methylation controls chromatin accessibility and nuclear factor binding at CNS2

To reveal the nature of this epigenetic mechanism, we first compared chromatin accessibility with ATAC-seq (Buenrostro, Giresi, Zaba, Chang, & Greenleaf, 2013) among CD4 Tn, Te, nT_reg_, and mock- and ASC-treated iT_reg_ cells. This experiment unbiasedly revealed significant differences between the chromatin accessibility of mock-treated iT_reg_ cells and that of ASC-treated iT_reg_ cells (Fig. 4*A, B*). At the *Foxp3* locus, we observed T_reg_-specific accessible chromatin at CNS2, in agreement with a published report (Samstein et al., 2012). Despite active Foxp3 transcription, CNS2 chromatin remained inaccessible in mock-treated iT_reg_ cells. Upon ASC treatment, CNS2 became accessible, suggesting that DNA demethylation drastically remodels CNS2 chromatin architecture.

**Fig. 4.**
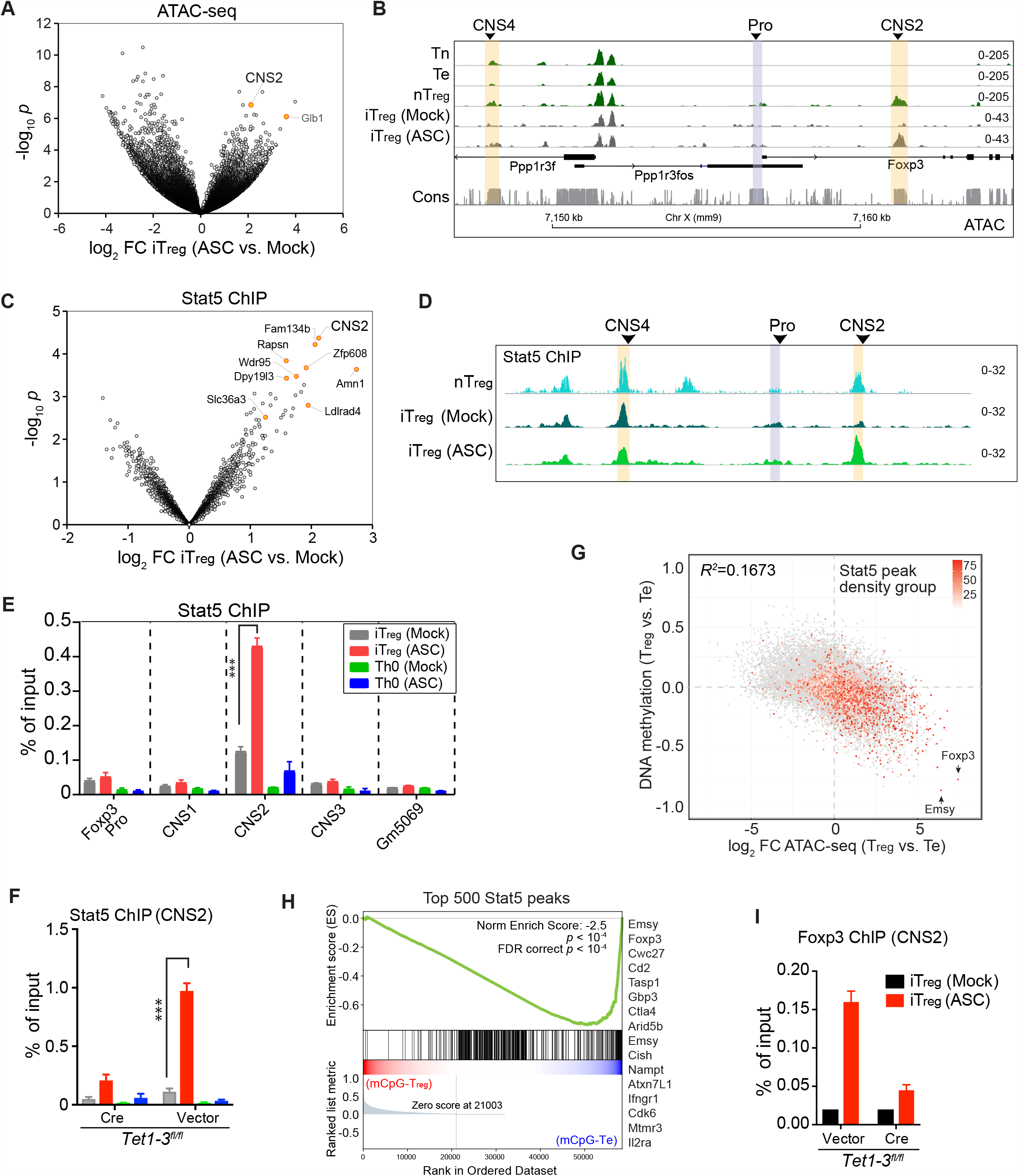
DNA methylation controls chromatin accessibility and nuclear factor binding at CNS2. (*A, B*) Genome-wide differential chromatin accessibility of mock- and ASC-treated iT_reg_ cells (*A*) and local chromatin accessibility around the *Foxp3* locus in nT_reg_ cells and in mock- and ASC-treated iT_reg_ cells derived from male *Foxp3*^*gfp*^ mice (*B*) assessed by ATAC-seq. Data represent >3 experiments. (*C, D*) Genome-wide differential Stat5 binding in mock- and ASC-treated iT_reg_ cells (*C*) and Stat5 binding sites around the *Foxp3* locus in nT_reg_ cells and in mock- and ASC-treated iT_reg_ cells (*D*). Cells were starved in IL-2–free media for 3 h and then stimulated with IL-2 (500 U/mL) for 30 min before being harvested for Stat5 ChIP-seq. Data represent 2 independent experiments. *(E)* Stat5 ChIP-qPCR with mock- and ASC-treated iT_reg_ and neutrally activated CD4 Tn cells (Th0 cells) after 30 min of IL-2 stimulation. Data represent 2 independent experiments. *(F)* Quantification of Stat5 binding at CNS2. *Tet1-3*^*fl/fl*^ CD4 Tn cells were transduced with Cre-expressing or control retrovirus 1 day after activation by TCR agonists and IL-2 in the presence or absence of TGF-β (i.e., iT_reg_ and Th0 culture conditions, respectively). ASC or PBS was added to media on day 3. Then, 3-4 days later, transduced cells were sorted by Thy1.1 reporter expression for Stat5 ChIP-qPCR. Data show means + SEMs of triplicates and represent 2 experiments. *(G)* Differential CpG methylation and chromatin accessibility of nT_reg_ and CD4 Te cells. Dots represent individual ATAC-seq peaks. Stat5-binding sites in nT_reg_ cells are superimposed. Heatmap shows the quantiles of Stat5 peak intensities. Data are merged from 2 replicates. *(H)* Gene set enrichment analysis of the relationship between the top 500 Stat5-binding sites and the differential DNA methylation of nT_reg_ and Te cells. Genes related to the top 16 Stat5 peaks are listed on the right. *(I)* Quantification of Foxp3 binding at CNS2 with ChIP-qPCR. Ablation of *Tet1-3* and iT_reg_ cell induction were performed as described in (*F*). See also Fig. S5.

Increased chromatin accessibility of CNS2 may facilitate nuclear factor binding, thus rewiring the regulatory circuits controlling Foxp3 transcription. To test this possibility, we assessed Stat5 binding at CNS2 before and after DNA demethylation because of the reported roles of IL-2/Stat5 signaling in T_reg_ cell induction and lineage stability (Chinen et al., 2016; Fan et al., 2018; Feng et al., 2014). Stat5 ChIP-seq revealed 3,286 Stat5-binding sites, among which 164 showed significantly different (*p*<0.05) intensities between mock- and ASC-treated iT_reg_ cells (Fig. 4*C*). Stat5 binding at CNS2 in mock-treated iT_reg_ cells was barely noticeable, and ASC treatment increased Stat5 binding at CNS2 to a level comparable to that in nT_reg_ cells (Fig. 4*D*), indicating a causal role of DNA demethylation. In comparison, Stat5 binding at enhancer CNS4 was independent of ASC treatment. Similar to CNS2, ASC treatment induced DNA demethylation, open chromatin, and enhanced Stat5 binding at many other regions (e.g., *Fam134b* and *Ldlrad4*) (Fig. S5*A, B*). In some cases, ASC treatment resulted in DNA hypomethylation and increased Stat5 binding without changing chromatin accessibility (Fig. S5*C*-*F*). Despite these differences, our data demonstrated an epigenetic switch induced by Tet/ASC– induced DNA demethylation that controls Stat5 binding.

To test whether increased Stat5 binding at CNS2 was T_reg_ cell–specific, we used ChIP-qPCR to assess Stat5 binding at *Foxp3 cis*-regulatory elements in mock- and ASC-treated Th0 (activated CD4 Tn cells in neutral conditions) and iT_reg_ cells. We observed Stat5 binding at CNS2 in ASC-treated iT_reg_ cells but not in mock-treated iT_reg_ cells or ASC-treated Th0 cells (Fig. 4*E*), in agreement with chromatin accessibility of CNS2 in ASC-treated iT_reg_ cells. To test whether Stat5 binding at CNS2 in ASC-treated iT_reg_ cells depends on Tet enzymes, we used retroviral Cre to acutely ablate *Tet1-3* in differentiating iT_reg_ cells two days before ASC treatment. This led to an abrupt reduction of Stat5 binding at CNS2 (Fig. 4*F*), confirming that the effect of ASC was achieved via Tet enzymes. Although the transition of Tet/ASC–induced DNA demethylation cannot be readily captured during T_reg_ cell development *in vivo*, Stat5 binding was highly enriched at hypomethylated, accessible chromatin regions in nT_reg_ cells (Fig. 4*G, H*), suggesting that DNA methylation controls chromatin accessibility and Stat5 binding, probably as a general mechanism.

DNA methylation may also control the binding of other nuclear factors at CNS2. To test this notion, we examined Foxp3 association with CNS2, a feedforward mechanism proposed to stabilize T_reg_ fate (Maruyama, Konkel, Zamarron, & Chen, 2011; Rudensky, 2011; Samstein et al., 2012). Indeed, we observed increased Foxp3 binding at CNS2 in ASC-treated *Tet1-3*^*fl/fl*^ iT_reg_ cells, which was diminished when cells were transduced with Cre-expressing retrovirus before their differentiation (Fig. 4*I*). Although the factors that differentially bind at hypomethylated CNS2 remain to be fully determined, DNA demethylation appears to reprogram the regulatory circuits via CNS2 to stabilize Foxp3 transcription by modulating the binding of nuclear factors.

### DNA demethylation stabilizes qualitative Foxp3 transcription *in vitro* in the absence of Foxp3 feedforward regulation

To reveal the roles of the regulatory circuits assembled via hypomethylated CNS2, we first assessed the contribution of IL-2/Stat5 signaling. Surprisingly, enhanced Stat5 binding at CNS2 appeared to not account for stable, qualitative Foxp3 expression in ASC-treated iT_reg_ cells, shown by the percentage of Foxp3^+^ cells after IL-2 deprivation (Fig. 2*C*). Consistently, low levels of IL-2 executed only a subtle effect on the stability of Foxp3 expression in WT nT_reg_ cells (Feng et al., 2014). Compared with the hyper instability of Foxp3 expression in *Tet2/Tet3*–deficient or mock-treated iT_reg_ cells, these results suggest a possibility that other factors bound at hypomethylated CNS2 might play a major role.

Next, we tested whether Foxp3 binding at hypomethylated CNS2 provides a strong feedforward mechanism to stabilize Foxp3 transcription, as similar circuits are widely used in cell fate determination (Urban & Johnston, 2018). To this end, we used *Foxp3*^*loxP-Thy1*.*1-Stop-loxP-gfp*^ (*Foxp3*^*LSL*^) knock-in mice (Hu et al, submitted) to report Foxp3 transcription by Thy1.1 expression without producing functional Foxp3 protein, thus they are comparable to *Foxp3*^*null*^ or scurfy mice (Fig. 5*A*). Because these mice have very few CD4 Tn cells as a result of severe immune activation, we generated chimeric mice with WT CD45.1 and *Foxp3*^*LSL*^ CD45.2 bone marrows mixed at a 1:1 ratio to suppress autoreactive conventional T cells (Fig. 5*B*). We isolated *Foxp3*^*LSL*^ CD4 Tn cells (CD45.2) from these healthy chimeric mice and induced Thy1.1-expressing “wannabe” iT_reg_ cells in T_reg_-induction conditions with or without supplemented ASC. We then sorted Thy1.1^+^ cells after induction to assess the stability of Thy1.1 expression in media containing JQ1, recombinant IL-2, or IL-2 neutralization antibodies.

**Fig. 5.**
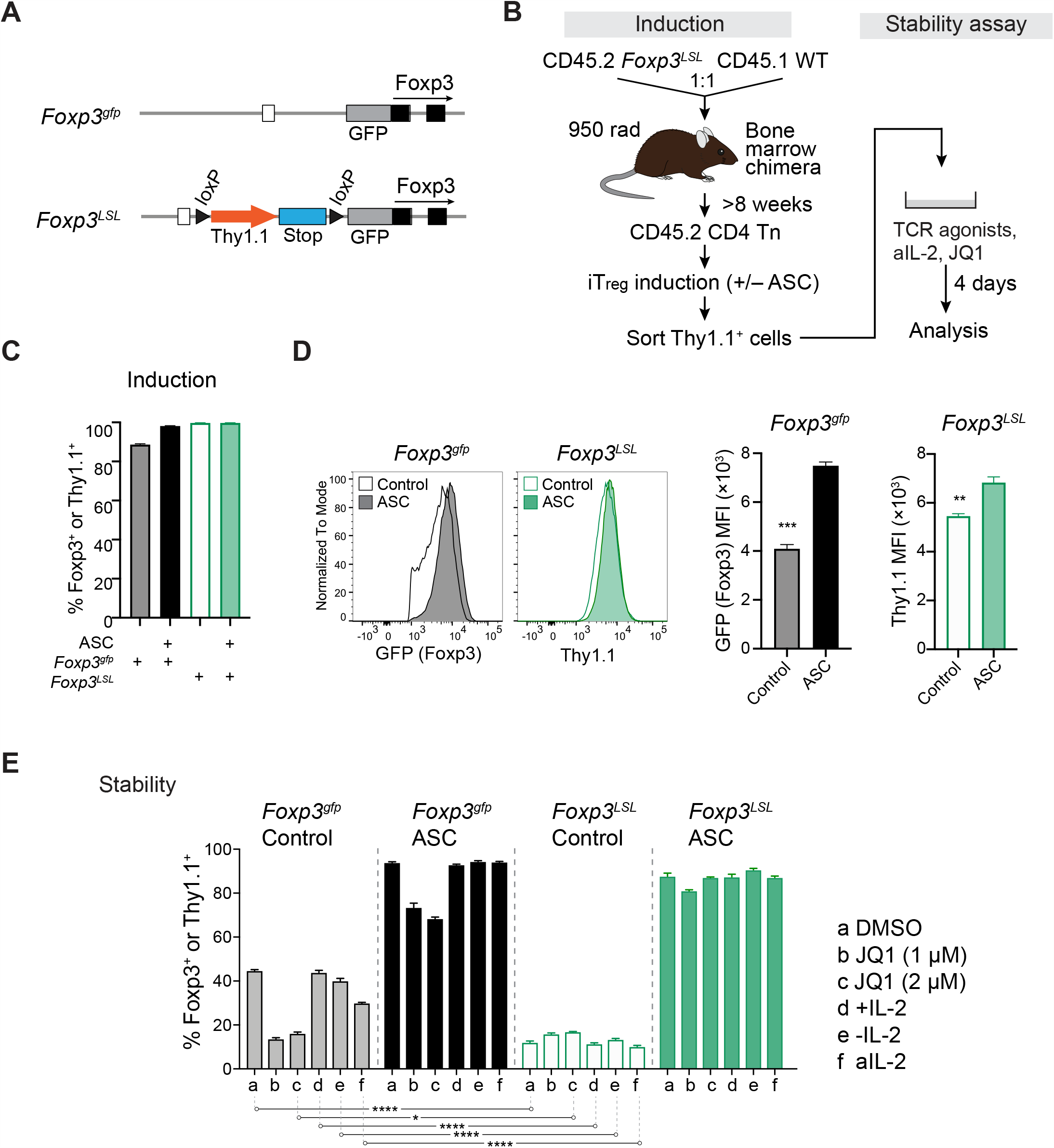
DNA demethylation stabilizes qualitative Foxp3 transcription *in vitro* in the absence of Foxp3 feedforward regulation. *(A)* Schematic of *Foxp3*^*gfp*^ and *Foxp3*^*LSL*^ knock-in mice. The latter reports Foxp3 transcription by Thy1.1 expression without producing Foxp3 protein in the absence of Cre recombinase. *(B)* Experimental procedures of Thy1.1 induction and stability assays *in vitro*. Mixed bone marrow chimeras of CD45.1 WT and CD45.2 *Foxp3*^*LSL*^ mice were generated. Thy1.1-expressing wannabe iT_reg_ cells were induced from *Foxp3*^*LSL*^ CD4 Tn cells with or without ASC treatment. These cells were FACS-sorted to assay the stability of Thy1.1 expression *in vitro*. Similarly, WT *Foxp3*^*gfp*^ CD4 Tn cells were used to examine the induction and stability of Foxp3 expression. (*C, D*) Induction of Foxp3 and Thy1.1 expression from *Foxp3*^*gfp*^ or *Foxp3*^*LSL*^ CD4 Tn cells, respectively, in the presence or absence of ASC. Data show means + SEMs of triplicates and represent 2 independent experiments. ** *p*<0.01, *** *p*<0.001 (unpaired, two-tailed *t* test). (*E*) Stability of Foxp3 expression in mock- and ASC-pretreated iT_reg_ cells and Thy1.1 expression in wannabe iT_reg_ cells. Sorted GFP^+^ (Foxp3^+^) or Thy1.1^+^ cells were restimulated with TCR agonists in media containing DMSO (100 U/mL IL-2), JQ1 (100 U/mL IL-2), IL-2 (100 U/mL; +IL-2), no recombinant IL-2 (-IL-2), or aIL-2 (25 µg/mL) for 4 days before analysis. Data show means + SEMs of triplicates and represent 2 independent experiments. * *p*<0.05, **** *p*<0.0001 (unpaired, two-tailed *t* test).

Lack of Foxp3 protein expression appeared to be dispensable for Thy1.1 induction efficiency (Fig. 5*C*), indicating that Foxp3 feedforward regulation plays a minor role in T_reg_ cell development. ASC treatment led to a mild increase of GFP-Foxp3 fusion protein in iT_reg_ cells and of Thy1.1 protein in wannabe iT_reg_ cells (Fig. 5*D*). In T_reg_ *in vitro* stability assay, more than 55% of control iT_reg_ cells and more than 80% of wannabe iT_reg_ cells failed to maintain Foxp3 or Thy1.1 expression, respectively, with the latter showing a significantly more instability (Fig. 5*E*). This result indicates that Foxp3 feedforward loop plays a role in sustaining Foxp3 transcription before Tet/ASC–induced DNA demethylation. After ASC treatment, however, lack of Foxp3 feedforward regulation did not affect the stability of Thy1.1 expression in the presence of JQ1 or upon IL-2 deprivation. As CNS2 is required for Tet/ASC– dependent stabilization of Foxp3 transcription, these data indicate that Stat5 and Foxp3 binding at hypomethylated CNS2 is insufficient to account for the drastically increased stability of Foxp3 transcription upon Tet/ASC–induced DNA demethylation.

### Foxp3 feedforward loop and maintenance of Foxp3 transcription and T_reg_ cell fitness *in vivo*

We then examined the roles of the Foxp3 feedforward loop *in vivo*. We sorted Thy1.1^+^ wannabe iT_reg_ cells after induction and cotransferred them with CD45.1 Tn and nT_reg_ cells into *Rag1*^*–/–*^ mice (Fig. 6*A*). Two weeks later, approximately 20% of control and 80% of ASC-pretreated wannabe iT_reg_ cells maintained Thy1.1 expression (Fig. 6*B, C*), further confirming that Tet-induced DNA demethylation stabilizes qualitative Foxp3 transcription in the absence of the Foxp3 feedforward loop. To test whether Foxp3 transcription can be stably maintained in the presence or absence of Foxp3 protein in nT_reg_ and wannabe T_reg_ cells developed *in vivo*, we sorted GFP^+^ CD4 T cells from male *Foxp3*^*gfp*^ mice and Thy1.1^+^ CD4 T cells from male *Foxp3*^*LSL*^ mice and co-transferred them with CD45.1 CD4 Tn and nT_reg_ cells into *Rag1*^*–/–*^ mice (Fig. 6*D*). Two weeks later, approximately 90% of nT_reg_ cells maintained Foxp3 expression and 70-80% of wannabe T_reg_ cells remained Thy1.1^+^ (Fig. 6*E*). The latter rate was comparable to that of ASC-pretreated wannabe iT_reg_ cells (Fig. 6*C*). Taken together, these results indicate that Foxp3 feedforward regulation is a minor contributor to Tet-induced stable qualitative Foxp3 transcription.

**Fig. 6.**
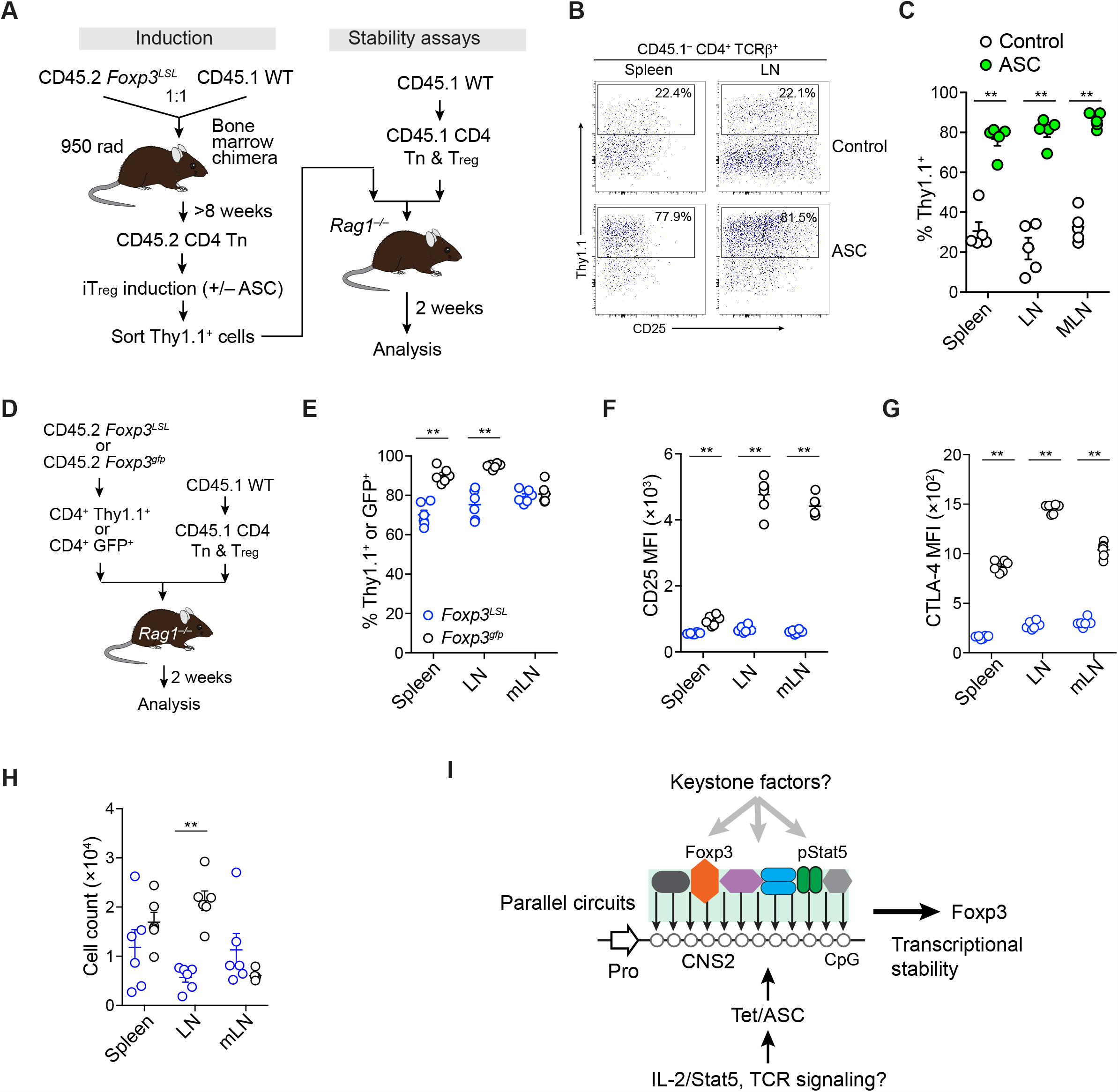
Roles of Foxp3 feedforward loop in maintaining Foxp3 transcription and T_reg_ cell fitness *in vivo*. (*A*) Experimental procedures for assessing the stability of Thy1.1 expression in wannabe iT_reg_ cells *in vivo*. CD4 Tn cells isolated from mixed bone marrow chimeric mice were used to induce Thy1.1 expression in T_reg_-induction media with or without supplemented ASC. Thy1.1^+^ cells were then sorted and co-transferred with CD45.1 CD4 T cells (including T_reg_ cells) into *Rag1*^*–/–*^ recipient mice. Cells were recovered 2 weeks later for analysis. (*B*-*C*) Thy1.1 expression in recovered CD45.1^−^CD4^+^TCRβ^+^ cells. n=5 each group. LN, lymph node; MLN, mesenteric lymph node. *** *p*<0.01 (Mann-Whitney tests). Data represent 2 experiments. (*D*-*H*) Stability of Foxp3 transcription in nT_reg_ or wannabe T_reg_ cells isolated from male *Foxp3*^*gfp*^ or *Foxp3*^*LSL*^ mice. FACS-sorted CD4^+^GFP^+^ or CD4^+^Thy1.1^+^ cells were co-transferred with CD45.1 CD4 T cells (including T_reg_ cells) into *Rag1*^*–/–*^ mice and recovered 2 weeks later to determine the percentages of GFP^+^ or Thy1.1^+^ cells among CD45.1^−^CD4^+^TCRβ^+^ cells (*E*). CD25 (*F*) and CTLA-4 (*G*) median fluorescence intensities were calculated in GFP^+^ and Thy1.1^+^ cells. GFP^+^ and Thy1.1^+^ cells were also counted in recovered cells (*H*). n=6 each group. ** *p*<0.01 (Mann-Whitney tests). (*I*) Model: During T_reg_ cell development, Tet enzymes demethylate the *Foxp3 cis*-regulatory elements, specifically 12 CpG sites within enhancer CNS2, to stabilize Foxp3 transcription. This process is accompanied by enhanced chromatin accessibility of CNS2 and association of CNS2 with Stat5 and Foxp3. However, the contribution of Stat5 and Foxp3 binding at CNS2 to the stable, qualitative Foxp3 expression is minor, suggesting that the parallel circuits established by CNS2 demethylation act in a partially redundant manner to maintain Foxp3 expression. Uncharacterized keystone factors in which single deficiency may severely impair the stability of qualitative Foxp3 transcription could also govern CNS2 function.

Despite comparable stability of qualitative Foxp3 transcription, absence of Foxp3 protein reduced CD25 and CTLA-4 expression in a cell intrinsic manner (Fig. 6*F, G*), consistent with the reported function of Foxp3 (Fontenot et al., 2003; M. A. Gavin et al., 2007; Hori et al., 2003). Because CD25 confers on IL-2 receptors an approximately 100-fold higher affinity to IL-2 (Malek & Castro, 2010) and IL-2 signaling plays a critical role in T_reg_ cell survival and fitness, lower levels of CD25 may contribute to the decrease in wannabe T_reg_ cell numbers (Fig. 6*H*). Thus, Stat5 and Foxp3 binding at hypomethylated CNS2 may play only a minor role in stabilizing qualitative Foxp3 transcription after Tet-induced DNA demethylation (Fig. 6*I*). Our data suggest that other factors and mechanisms may contribute to the abrupt switch of the driving force that stabilizes qualitative Foxp3 transcription upon Tet-dependent DNA demethylation during T_reg_ cell development.

## DISCUSSION

We aim to uncover the nature of qualitative and quantitative regulations of Foxp3 transcription during the transition from T_reg_ cell induction to lineage maintenance. Tet enzymes were proposed to stabilize qualitative Foxp3 transcription in *cis* by demethylating the *Foxp3* enhancer CNS2. However, this model has not been directly tested, leaving its real role and underlying mechanisms uncertain. Given the potential global effect of Tet-dependent DNA demethylation, it is unclear whether and to what extent Tet enzymes stabilize Foxp3 expression through CNS2. Besides observed DNA hypomethylation of CNS2 in T_reg_ cells and impaired stability of qualitative Foxp3 expression in Tet- or CNS2-deficient T_reg_ cells (Sakaguchi et al., 2020), how DNA demethylation stabilizes Foxp3 expression in its native genomic context is also unclear.

As no markers are available to readily distinguish the developmental stages of T_reg_ cells *in vivo*, we modeled T_reg_ lineage commitment using *ex vivo*–isolated mouse T cells. Limited ASC in regular culture medium enables us to control Tet enzymatic activity via supplemented ASC (Sasidharan Nair et al., 2016; Yue et al., 2016). This treatment reliably demethylates *Foxp3 cis*-regulatory elements and stabilizes qualitative Foxp3 transcription in iT_reg_ cells, recapitulating the key features of T_reg_ lineage commitment (Rudensky, 2011; Sakaguchi et al., 2020). Using this model, we directly assessed the role of Tet-dependent DNA demethylation in stabilizing Foxp3 expression, thereby affecting T_reg_ lineage identity in *cis* via enhancer CNS2. ASC-treated iT_reg_ cells and nT_reg_ cells share similar DNA methylation levels, chromatin accessibility, and Foxp3 and Stat5 binding at CNS2. However, nT_reg_ and ASC-treated iT_reg_ cells are not identical in DNA methylation patterns, indicating considerable differences in Tet recruitment and/or enzymatic activity. Nonetheless, ASC treatment stabilizes Foxp3 transcription in iT_reg_ cells in a CNS2-dependent manner, demonstrating the essential role of the Tet/ASC-CNS2 axis in sustaining qualitative Foxp3 expression and T_reg_ lineage identity. It also distinguishes the *cis* and *trans* effects of Tet/ASC-mediated DNA demethylation on T_reg_ lineage stability.

Tet/ASC–induced DNA demethylation increased the chromatin accessibility of CNS2, accompanied by markedly enhanced Stat5 and Foxp3 binding. These observations provide a mechanistic understanding of how DNA methylation controls the stage-specific activity of CNS2. Although similar phenomena have been shown by *in vitro* binding and reporter assays (H. P. Kim & Leonard, 2007; Polansky et al., 2010; Zheng et al., 2010), in the absence of native genomic context, such results did not reveal the exact function of the factors bound at hypomethylated CNS2. We found that demethylation of CNS2 induces a drastic switch of the regulatory modes of Foxp3 transcription, enabling stable, qualitative Foxp3 expression in adverse conditions while maintaining its quantitative regulation by IL-2 and permissive histone acetylation signaling. Counterintuitively, however, enhanced Stat5 binding at hypomethylated CNS2 appears to play only a minor role in stabilizing Foxp3 expression. Similarly, ASC treatment largely stabilizes Foxp3 transcription in the absence of the Foxp3 feedforward loop with or without IL-2 deprivation. Therefore, the real functions of the factors bound at hypomethylated CNS2 should be examined experimentally in the native genomic context during T_reg_ lineage maintenance.

Our study of Stat5 and Foxp3 binding raised the question of whether a single factor bound at hypomethylated CNS2 accounts for the drastically enhanced stability of qualitative Foxp3 transcription. Previous studies identified several factors bound at the *Foxp3 cis*-regulatory elements to maintain Foxp3 expression, such as CBFβ (Rudra et al., 2009). However, it is unclear to what extent their roles are mediated by CNS2’s demethylation. The 12 CpG sites scattered in the CNS2 region undergo a drastic demethylation during T_reg_ lineage commitment in a Tet/ASC–dependent manner, suggesting that DNA demethylation could control the binding of multiple nuclear factors at CNS2 (Fig. 6*I*). If each of them contributes a part to CNS2’s function, then CNS2 demethylation likely creates “parallel” regulatory circuits to stabilize Foxp3 transcription, so loss of any individual factor’s binding would not generate a catastrophic effect. Therefore, T_reg_ lineage identity could be maintained by these partially redundant mechanisms to oppose the fluctuations of individual proteins caused by adverse environments or genetic variations. In addition, particular proteins or keystone factors associated with hypomethylated CNS2 alone may be essential for stable Foxp3 expression, and their single deficiency could severely impair the stability of Foxp3 expression. These proteins may be the unique constituents of CNS2 or indispensable components of enhanceosomes in general. Given the exceptional stability of T_reg_ lineage identity, the keystone factors are likely abundant in T_reg_ cells, akin to housekeeping genes or generic transcription factors, such that robust Foxp3 transcription can be sustained in the presence of various perturbations. Therefore, identifying and characterizing the proteins associated with hypomethylated CNS2 will produce important new insights into the mechanisms governing T_reg_ lineage stability.

In summary, we modeled the processes of T_reg_ lineage commitment with mouse primary T cells to directly investigate the role and underlying mechanisms of Tet/ASC–induced DNA demethylation in controlling qualitative Foxp3 transcription and, thereby, T_reg_ lineage identity. Our results proved the proposed model of Tet/ASC-CNS2 axis in sustaining Foxp3 expression, distinguished it from alternative mechanisms, and uncovered an epigenetic switch of the driving force of qualitative Foxp3 expression after Tet-dependent DNA demethylation during T_reg_ cell development. Thus, our study consolidates the basis for further investigation of the mechanisms controlling T_reg_ lineage commitment and maintenance, an essential step to fully understanding T_reg_-mediated immune tolerance in various immunological settings.

## ACKNOWLEDGMENTS AND FUNDING SOURCES

We thank Alexander Rudensky for providing the mouse strains, the Hartwell Center for high-throughput sequencing, the Flow Cytometry Core facility for assistance with cell sorting and analysis, Jaeki Min and Zoran Rankovic for providing epigenetic inhibitors, Cherise Guess for editing the manuscript, and Ruopeng Feng for help with the Cut&Run experiments. X.Z. was supported by an APO Special Postdoctoral Fellowship of St. Jude Children’s Research Hospital. This study was supported by the American Lebanese Syrian Associated Charities and the NIH grant R21AI146614 (Y. Feng.). The content is solely the responsibility of the authors and does not necessarily represent the official views of the National Institutes of Health.

## AUTHOR CONTRIBUTIONS

J.L. and Y. Feng. conceived of and designed the experiments and interpreted the results. X.Z. quantified nascent Foxp3 mRNA and provided critical comments. M.H. performed the Cut&Run experiments and interpreted the results. B.X. and Y. Fan analyzed the high-throughput sequencing data. R.C. provided assistance with FACS sorting. J.H. provided *Tet1-3*^*fl/fl*^ mice. J.L. and Y. Feng. wrote the manuscript.

## CONTACT FOR REAGENT AND RESOURCE SHARING

Further information and requests for resources and reagents should be directed to Yongqiang Feng.

## SUPPLEMENTARY INFORMATION

**Fig. S1.**
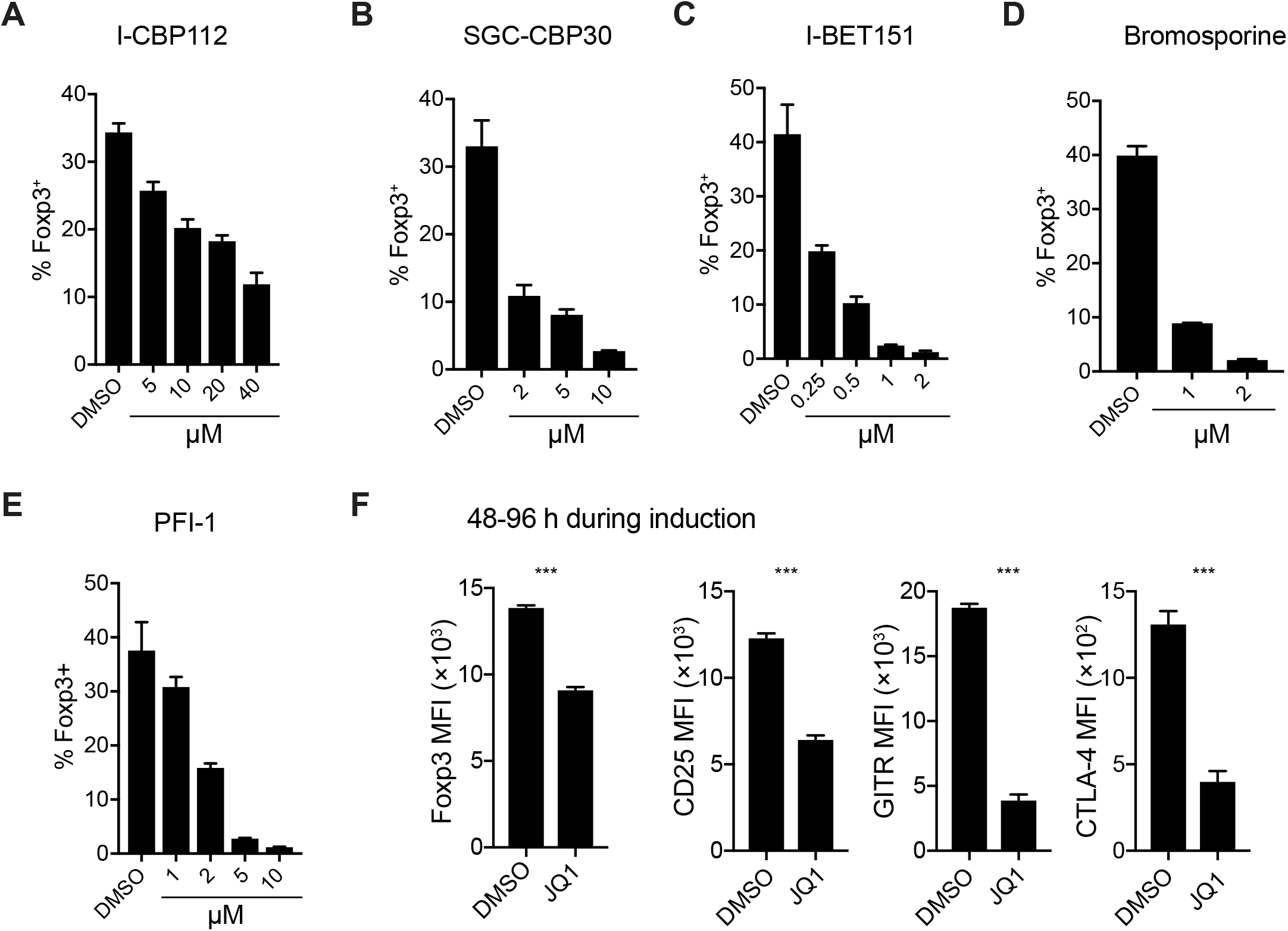
Histone acetylation drives Foxp3 induction during T_reg_ cell development *in vitro*. (*A*-*E*) Titrated amounts of I-CBP112 (*A*), SGC-CBP30 (*B*), I-BET151 (*C*), bromosporine (*D*), or PFI-1 *(E)* were added in culture media during T_reg_ cell induction *in vitro*. Cells were harvested on day 4 to analyze Foxp3 expression. *(F)*Effect of JQ1, when added from day 3 to day 4 during T_reg_ cell induction, on the median fluorescence intensity of Foxp3, CD25, GITR, and CTLA-4 in CD4^+^Foxp3^+^ cells. Data show means + SEMs of triplicates and represent 2 independent experiments. \*\*\**p*<0.001 (unpaired, two-tailed *t* test). Related to Fig. 1.

**Fig. S2.**
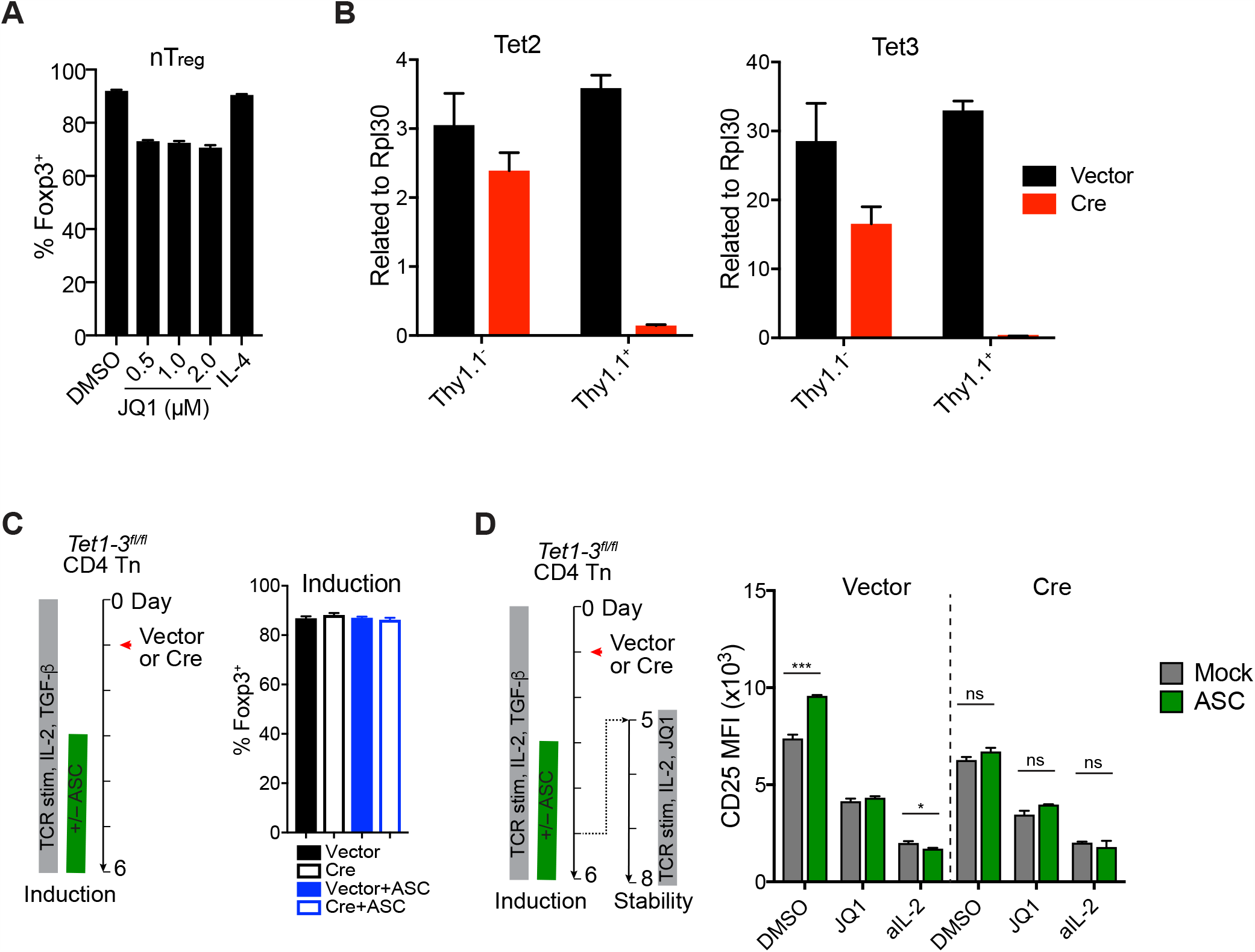
Effect of JQ1 on the stability of Foxp3 expression in nT_reg_ and iT_reg_ cells. *(A)*The nT_reg_ cells doubled-sorted from *Foxp3*^*gfp*^ mice were stimulated with beads coated with anti-CD3 and anti-CD28 antibodies in the presence of 500 U/mL recombinant IL-2 and either DMSO, IL-4, or titrated amounts of JQ1 for 4 days. Live cells were gated to analyze Foxp3 expression. Data show means + SEMs of triplicates and represent >2 independent experiments. \*\*\**p*<0.001 (unpaired, two-tailed *t* tests). *(B)*Quantification of *Tet2* and *Tet3* deletion. CD4 Tn cells isolated from *Tet1-3*^*fl/fl*^ mice were cultured in T_reg_-induction conditions. Cells were then transduced with vector or Cre-expressing retrovirus (reported by Thy1.1 expression); 3-4 days later, Thy1.1^+^ and Thy.1^−^ cells were sorted for genomic DNA extraction and qPCR. Data show means + SEMs of triplicates. *(C)*Effect of *Tet*-deficiency and ASC treatment on Foxp3 induction efficiency. Data show means + SEMs of triplicates and represent 2 independent experiments. *(D)*Effect of *Tet*-deficiency and ASC treatment on CD25 expression in cells that maintained Foxp3 expression during T_reg_ stability assay. Data show means + SEMs of triplicates and represent >2 independent experiments. ns, not significant, \**p*<0.05, \*\*\**p*<0.001 (unpaired, two-tailed *t* test). Related to Fig. 2.

**Fig. S3.**
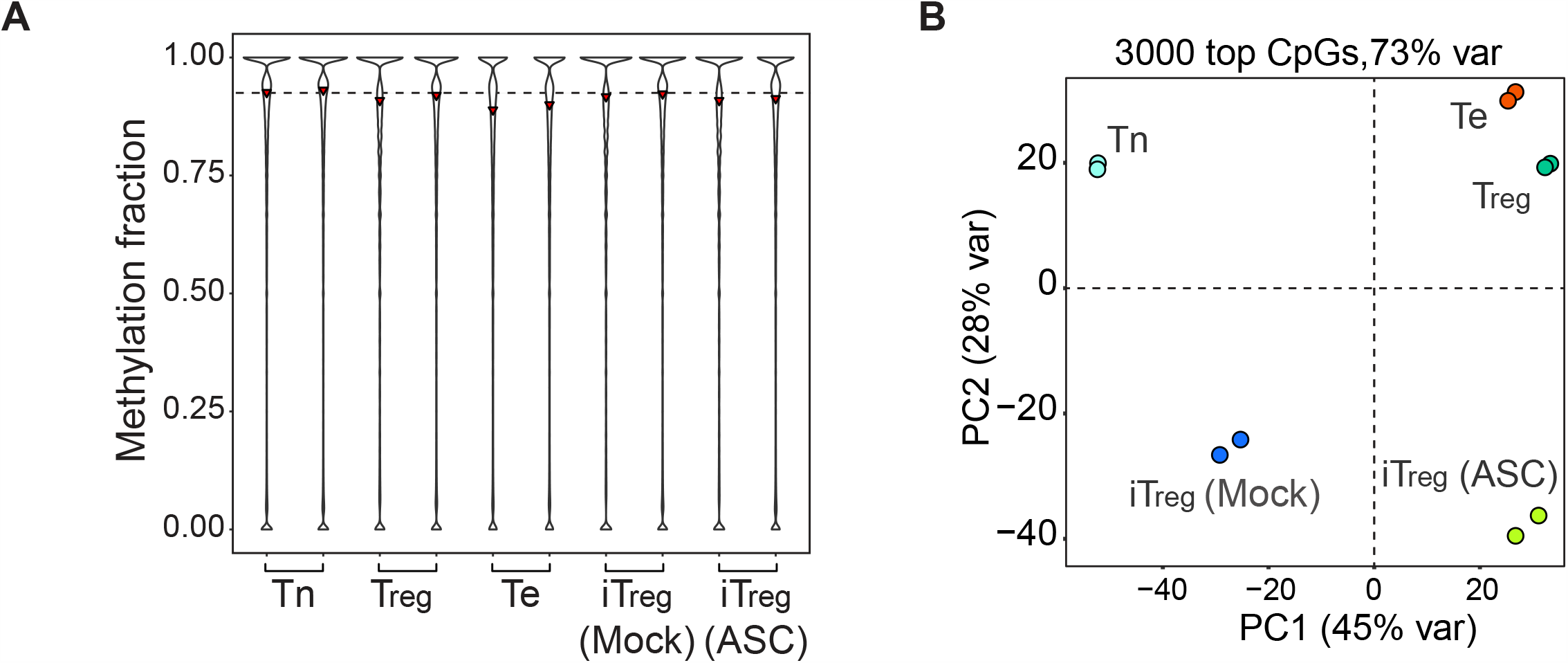
Whole-genome bisulfite sequencing of iT_reg_ cells treated with ASC. *(A)*Overall methylation fraction of CpG sites in CD4 Tn, nT_reg_, Te, and mock- and ASC-treated iT_reg_ cells. Whole-genome bisulfite sequencing was performed by using cells sorted from male *Foxp3*^*gfp*^ mice with or without being cultured *in vitro*. Data were merged from 2 biological replicates. *(B)*Principal component analysis and clustering of the top 3000 most-variable CpG sites, explaining 73% of total variations (var). Related to Fig. 3

**Fig. S4.**
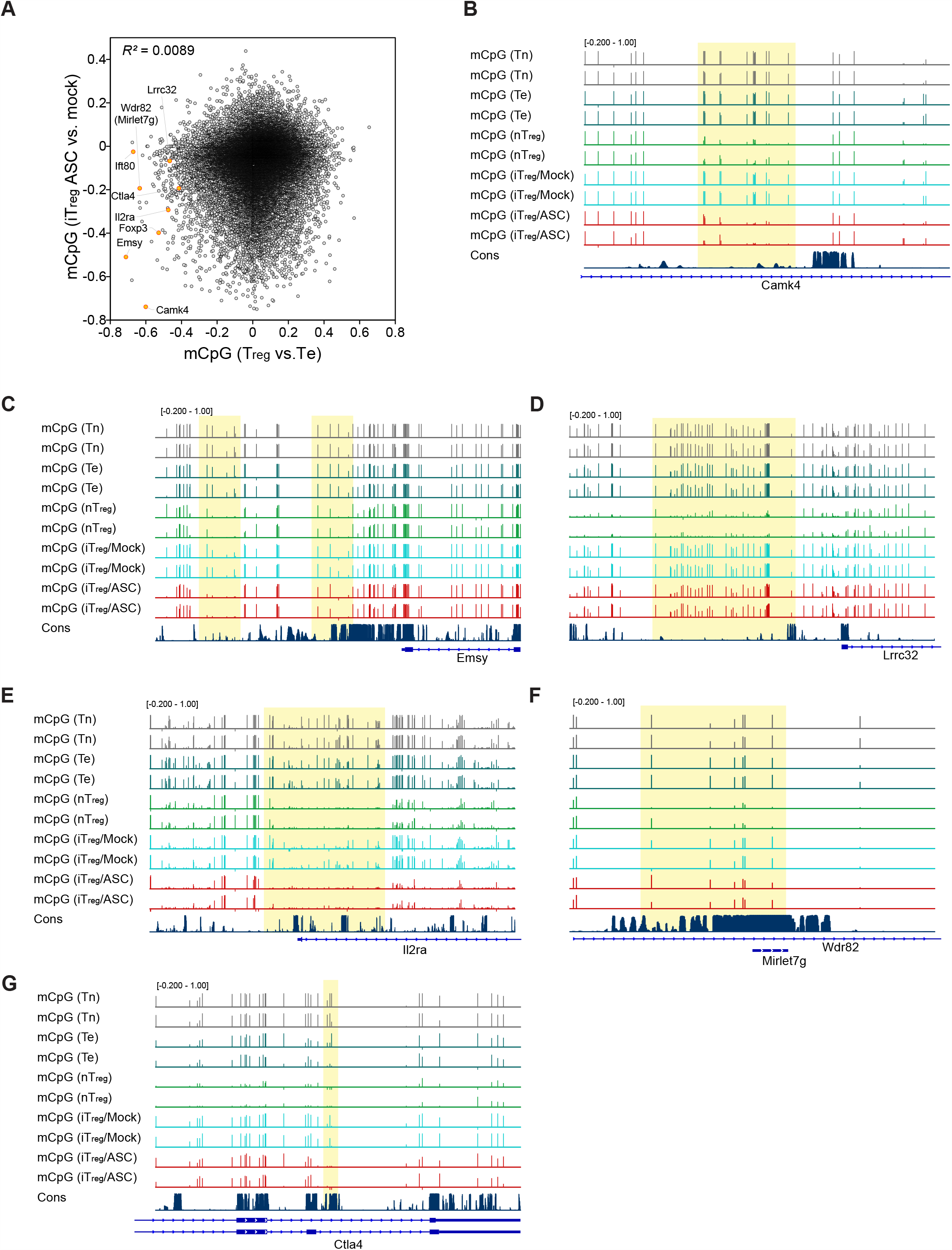
Representative regions subjected to Tet/ASC–dependent DNA demethylation. (*A*) Differentially methylated CpG sites in ASC-treated iT_reg_ and nT_reg_ cells. Differences in mCpG levels of nT_reg_ and Te cells were plotted against those of mock-0061nd ASC-treated iT_reg_ cells. WGBS with 40× coverage was performed by using cells derived from male *Foxp3*^*gfp*^ mice. Data were merged from 2 biological replicates. (*B*-*G*) Representative regions (highlighted) showing differences in mCpG levels of mock- and ASC-treated iT_reg_ cells. Vertical bars represent individual CpG sites; their heights indicate methylation levels from 0 to 1. All samples were plotted at the same scale. Data were merged from 2 biological replicates. Related to Fig. 4.

**Fig. S5.**
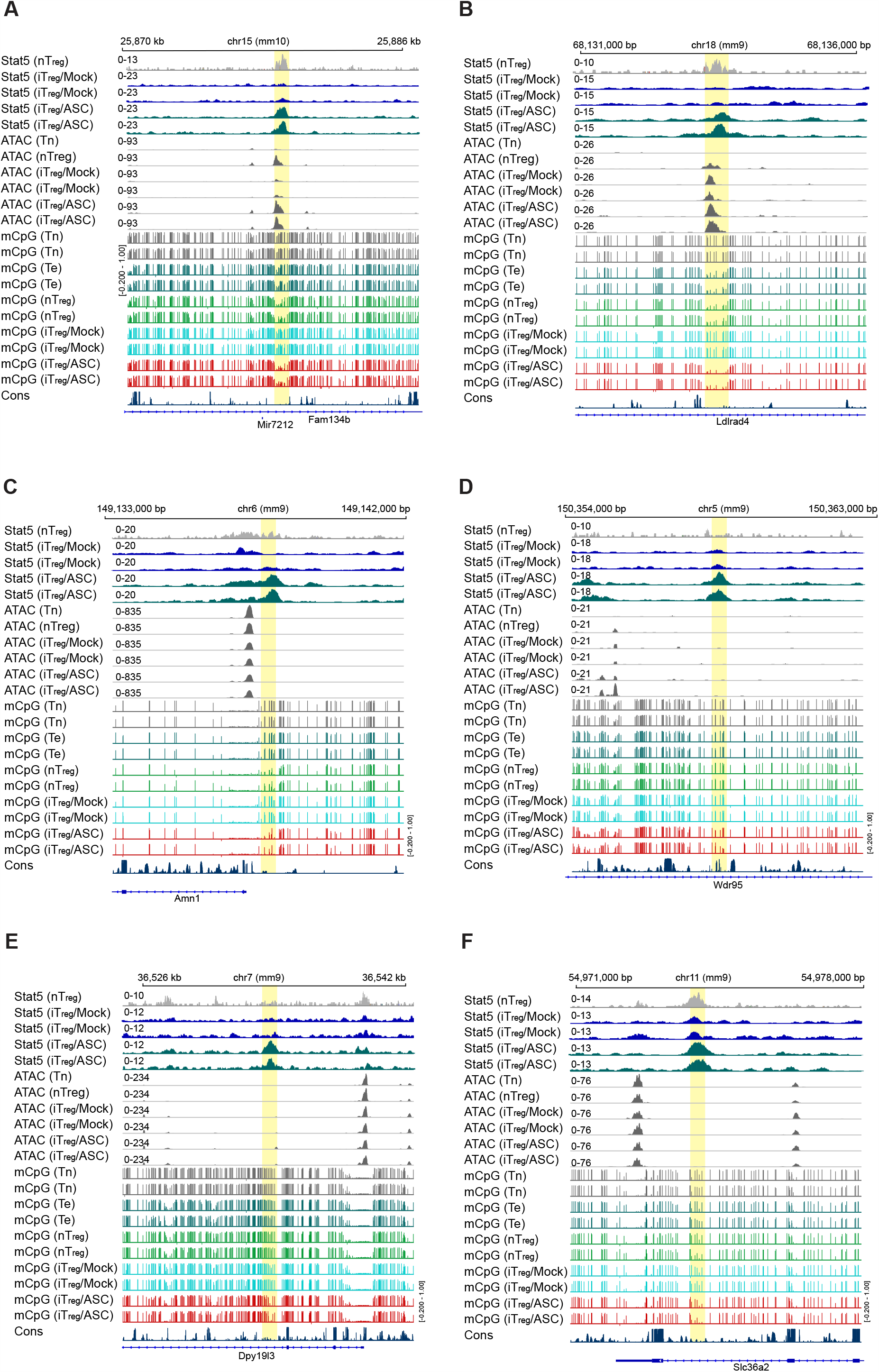
Representative regions with differential CpG methylation and Stat5 binding. *(A)* ATAC-seq, mCpG levels, and Stat5 ChIP-seq of nT_reg_ and mock- and ASC-treated iT_reg_ cells were aligned with DNA sequence conservation. Regions undergoing differential DNA methylation and Stat5 binding are highlighted. Related to Fig. 4.

**Supplementary Table 1.**
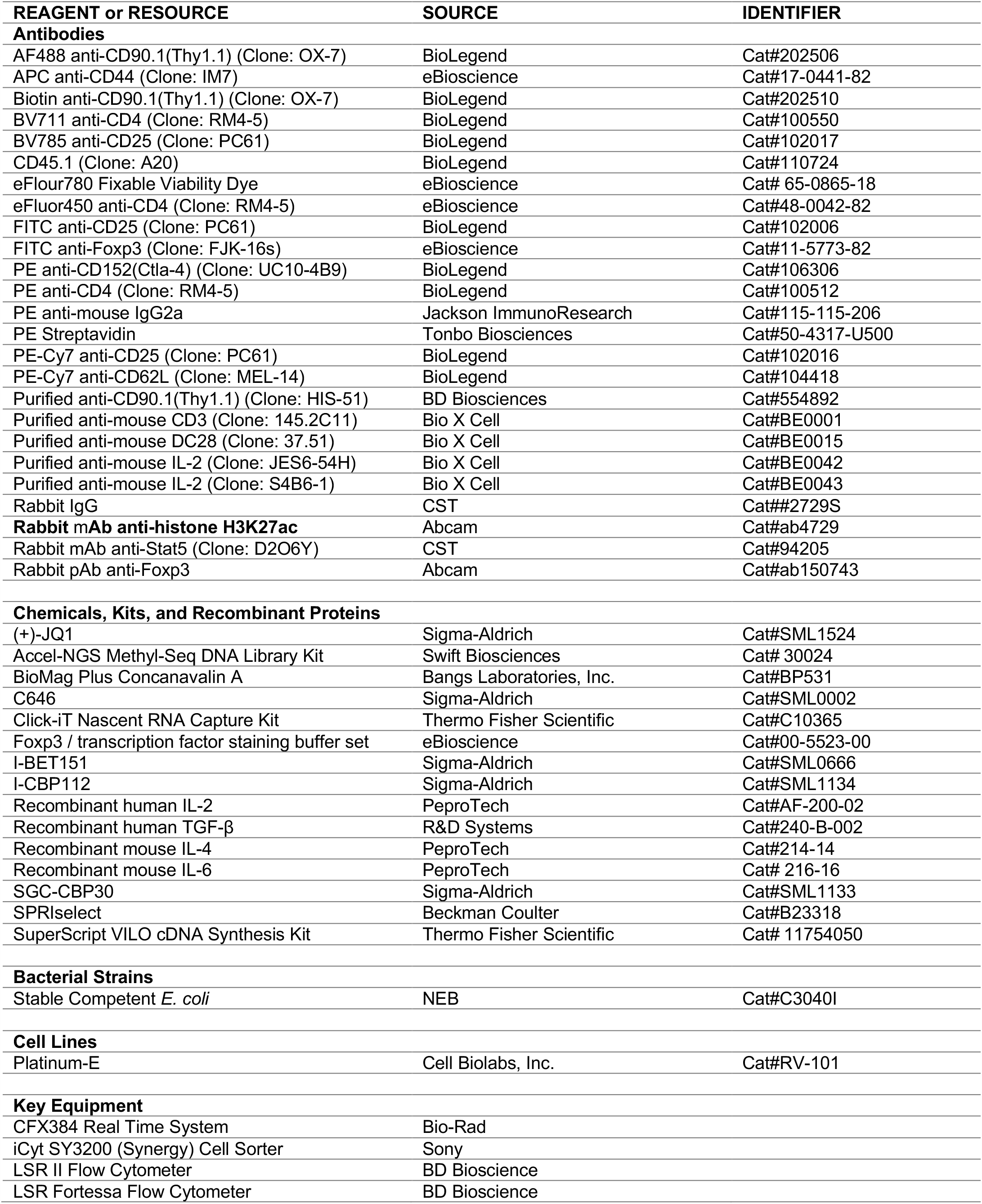
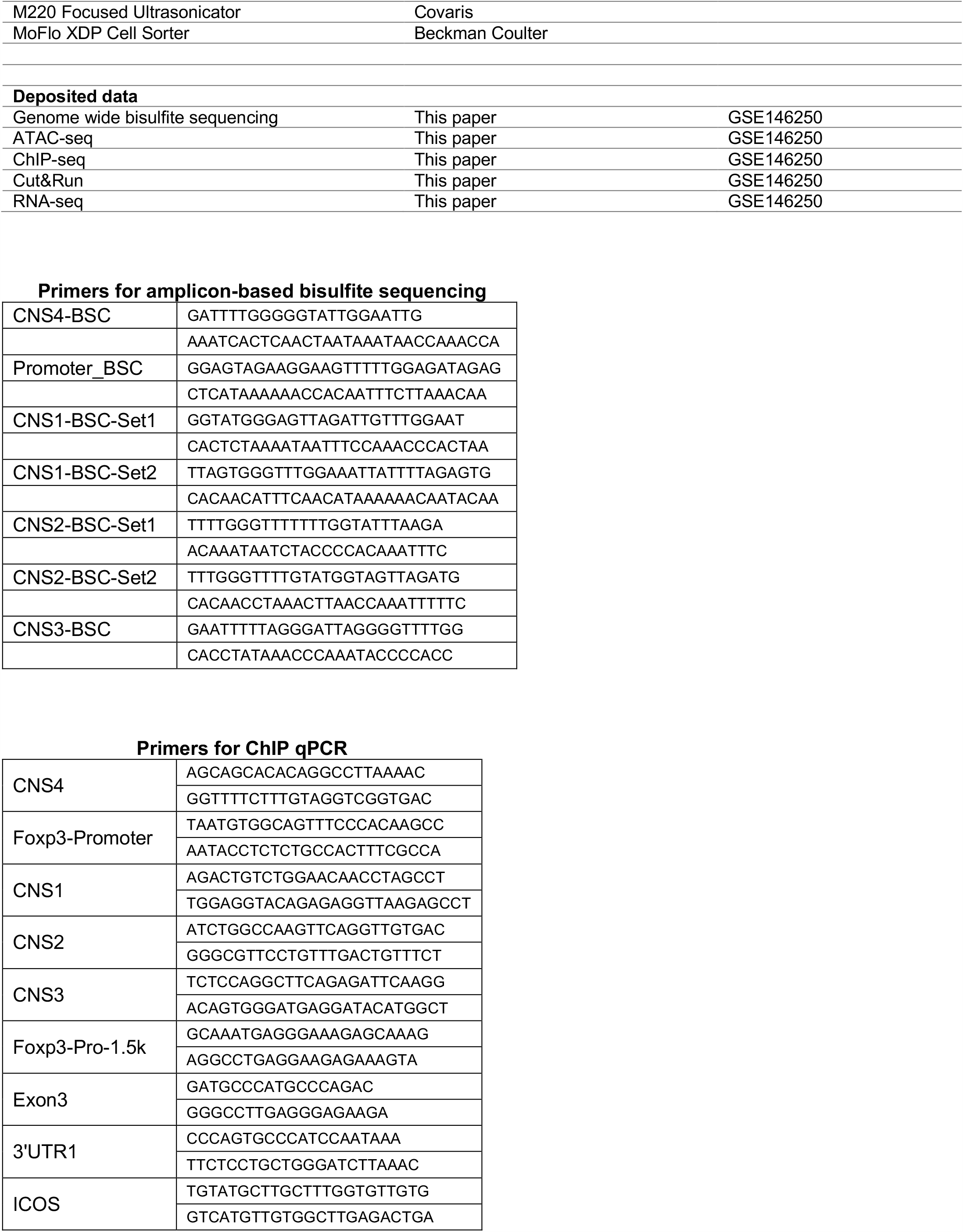
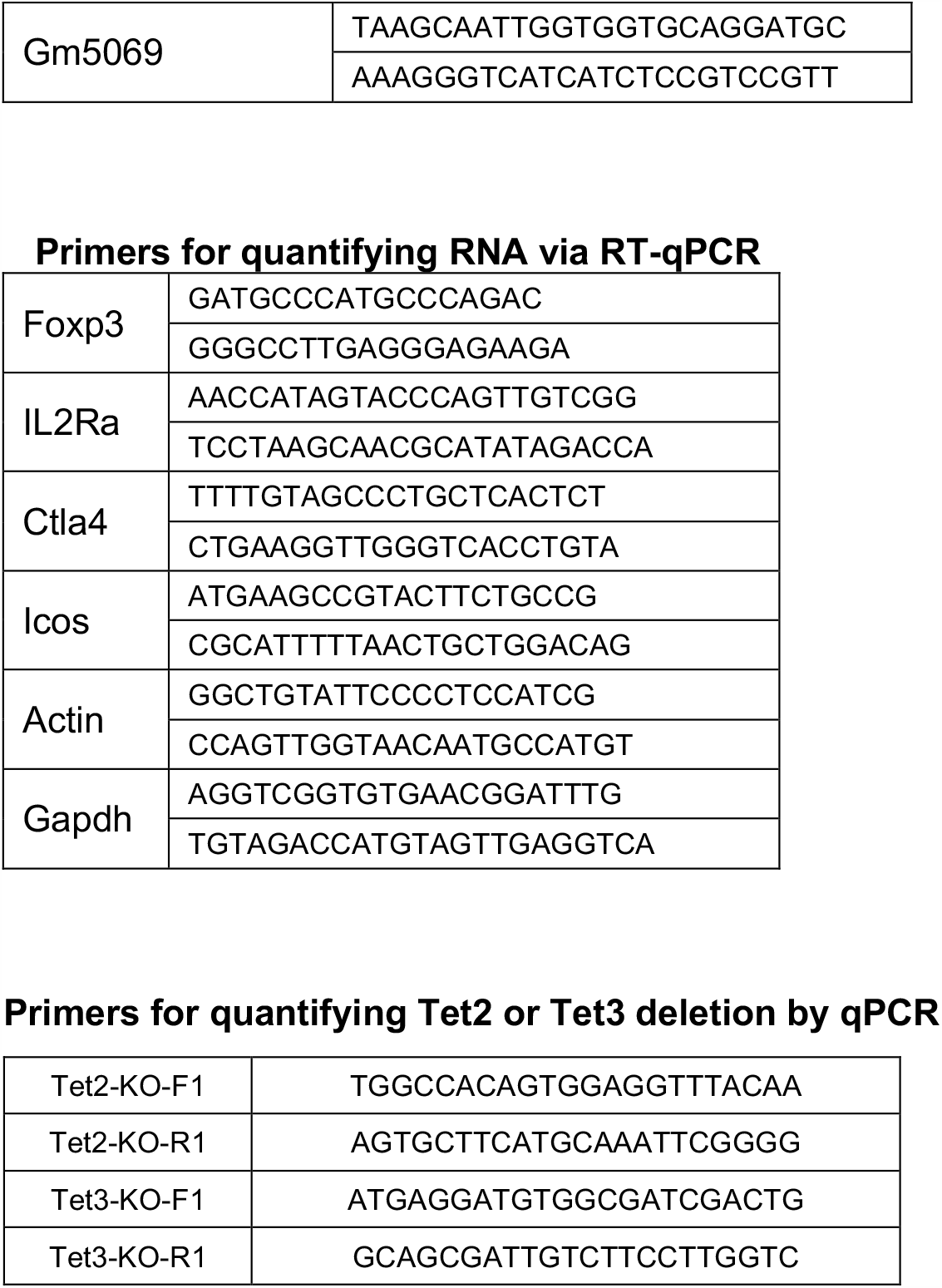
Key resources used in this study.

## MATERIALS AND METHODS

### Mice

All animal experiments were approved by St. Jude Children’s Research Hospital’s Institutional Animal Care and Use Committee (approval number 612). All mice were maintained and bred in a specific pathogen–free facility. Six-to twelve-week-old male and female mice were used for T_reg_ experiments. Because the *Foxp3* gene is on the X-chromosome, which undergoes random inactivation in females, male mice were used for epigenetic experiments, including whole-genome bisulfite sequencing, chromatin accessibility, ChIP, and Cut&Run assays. *Foxp3*^*gfp*^, *Foxp3*^*ΔCNS2-gfp*^, and *Tet1*^*fl/fl*^ *Tet2*^*fl/fl*^ *Tet3*^*fl/fl*^ (*Tet1-3*^*fl/fl*^) mice were described previously (Fontenot et al., 2005; Herzig et al., 2017; Zheng et al., 2010). *Foxp3*^*loxP-Thy1*.*1-Stop-loxP-gfp*^ (*Foxp3*^*LSL*^) mice were generated by inserting the loxP-Thy1.1-Stop-loxP-gfp cassette into a *Foxp3* intron before the coding sequence (Hu et al., submitted).

### T cell isolation and culture

CD4 T cells from the lymph nodes and spleens were enriched via EasySep Mouse CD4 T Cell Isolation Kits (STEMELL). CD4 naïve T cells (Tn, CD4^+^CD25^−^CD44^−^CD62L^hi^) and natural T_reg_ cells (CD4^+^GFP^+^) T cells were further sorted by FACS. T cells were cultured at 37°C, 5% CO_2_ in complete RPMI1640 medium (RPMI1640 supplemented with 10% fetal bovine serum [FBS], 2 mM L-glutamine, 1 mM sodium pyruvate, 1% non-essential amino acid, 10 mM HEPES, 20 µM 2-mercaptoethanol, 100 U/mL penicillin, and 100 mg/mL streptomycin) and indicated cytokines and compounds.

### T_reg_ cell induction and stability assays

Induction of T_reg_ or wannabe T_reg_ differentiation *in vitro* was conducted according to published protocols (Feng et al., 2015; Yue et al., 2016). Briefly, plates or dishes were precoated with anti-CD3 and anti-CD28 antibodies (1 µg/mL) in PBS (Supplementary Table 1) at 37°C for 2 h to culture FACS-sorted CD4 Tn cells from WT or *Foxp3*^*LSL*^ mixed bone marrow chimeras in complete RPMI1640 supplemented with recombinant human IL-2 (100 U/mL) and recombinant human TGF-β (1 ng/mL) for 4 days with or without 0.25 mM ascorbic acid-2-phosphate or indicated compounds. To test the stability of Foxp3 transcription, iT_reg_ or wannabe iT_reg_ cells that were sorted based on GFP-Foxp3 or Thy1.1 expression, respectively, after 4 days of induction were further cultured on plates pre-coated with anti-CD3 and anti-CD28 antibodies (1 µg/mL each), recombinant human IL-2 (100 U/mL), IL-2 neutralization antibodies (25 µg/mL each of JES6-54H and S4B6-1), or indicated cytokines or compounds for 4 more days. The nT_reg_ cells were cultured in the presence of Dynabeads Mouse T-Activator CD3/CD28 (Invitrogen) with 500 U/mL recombinant human IL-2 for 4 days and then given fresh media with recombinant human IL-2 and indicated cytokines or compounds for 4 more days. CD4 Tn cells were cultured on anti-CD3 and anti-CD28 antibody–coated plates or dishes with recombinant human IL-2 for 4 days to generate Th0 cells. After T_reg_ induction or stability assays, cells were collected and stained with viability dye and fluorophore-conjugated antibodies for flow-cytometry analysis.

### Flow-cytometry analysis

Cell staining and flow-cytometry analyses were performed as we previously described (Feng et al., 2014; Feng et al., 2015). Briefly, fluorophore-conjugated antibodies were used to stain cells (Supplementary Table 1). Cells were first stained with fixable viability dye, then incubated with indicated antibodies against cell surface markers followed by fixation/permeabilization with the Foxp3/Transcription Factor Staining Buffer Set (eBioscience) and intracellular staining for Foxp3, if needed. Cells were fixed in 1% paraformaldehyde for 10 min after staining. FACS analyses were performed on LSRII or LSR Fortessa (BD Biosciences) flow cytometers; data were analyzed via FlowJo (BD Biosciences).

### Retroviral packaging and transduction

Retroviral packing and transduction of mouse primary T cells were conducted according to our published protocols (Feng et al., 2015). Briefly, Platinum-E (Plat-E) cells were used to package the retrovirus. Puromycin (1 µg/mL) and blasticidin (10 µg/mL) were added to the culture media to maintain Plat-E cells according to the manufacturer’s manual (Cell Biolabs); 24 h before transfection, cells were seeded on new dishes in media without puromycin and blasticidin. Then, pMigR1-IRES-Thy1.1 or pMigR1-Cre-IRES-Thy1.1 plasmid (Feng et al., 2015) and pCl-Eco were cotransfected into these cells with TransIT-293 Transfection Reagent (Mirus); 2-3 days after transfection, viral supernatant was collected and filtered with 0.45-µm syringe filters, aliquoted, and stored at −80°C.

### Bisulfite sequencing

Cells used for bisulfite sequencing were sorted from male *Foxp3*^*gfp*^ mice with or without *in vitro* culture. The following markers were used to sort cells: for Tn, CD4^+^GFP^−^CD25^−^CD44^lo^CD62L^hi^; for Te, CD4^+^GFP^−^CD44^hi^CD62L^lo^; and for T_reg_, CD4^+^GFP^+^. Genomic DNA was prepared from sorted cells by proteinase K digestion followed by phenol:chloroform:isoamyl alcohol extraction and 2-propanol precipitation. More than 100 ng of genomic DNA in each sample was converted via EpiTect Bisulfite Kits (Qiagen). Bisulfite conversion efficiency was tested via PCR and Sanger sequencing of *Foxp3* enhancer CNS2 as we previously described(Feng et al., 2014). To perform whole-genome bisulfite sequencing, libraries were prepared from converted DNA by using the Accel-NGS Methyl-Seq DNA Library Kit (Swift Biosciences), analyzed for insert size distribution on a 2100 BioAnalyzer (Agilent Technologies,), and quantified by using the Quant-iT PicoGreen ds DNA assay (Life Technologies). Paired-end, 100-cycle sequencing was performed on a Hi-seq 4000 (Illumina) to achieve an average 40× coverage.

Bisulfite sequencing data were aligned to mouse genome mm9 by using BSMAP2.74 (Xi & Li, 2009). The methylation ratio for each CpG site was extracted by methratio.py from BSMAP2.74. The methylation ratio was then converted to bw file for visualization. The M values for the 3,000 most variable CpG sites were used for hierarchical clustering and PCA analyses. M value = log_2_ (C count/T count) at a CpG site (an offset value of 0.5 was added).

### ATAC Sequencing

ATAC-Seq was performed as previously reported (Buenrostro, Wu, Chang, & Greenleaf, 2015). Briefly, 5×10^4^ cells were FACS sorted and lysed in 300 µL of ice-cold lysis buffer (10 mM Tris-HCl, pH 7.5, 10 mM NaCl, 3 mM MgCl_2_, 0.1% NP-40). After centrifugation, the supernatant was removed; 50 µL of reaction mix containing 25 µL of TD buffer, 2.5 µL of TDE1 (Illumina Nextera DNA Library Preparation Kit), and 22.5 µL of nuclease-free water was immediately added to perform transposition at 42°C for 40 min. DNA was purified by using the NucleoSpin Gel and PCR Clean-up kit (MACHEREY-NAGEL). The transposed DNA was amplified by PCR for 10-12 cycles by using the Nextera DNA Library Preparation Kit and Nextera XT Indexing Kit (Illumina). The library DNA within the 150-to 500-bp range was enriched by AMPure XP beads (Beckman Coulter), quantified by NEBNext Library Quant Kit (NEB), and sequenced with paired-end 100-cycles on a HiSeq 4000 sequencer (Illumina). Paired-end reads of 100bp were trimmed by cutadapt (version 1.9, paired-end mode, default parameter with “-m 25 −O 6”) (Martin, 2011) and aligned to mouse genome mm9 (MGSCv37 from Sanger) by BWA (version 0.7.12-r1039, default parameter)(H. Li & Durbin, 2009). Duplicate reads were marked by biobambam2 (version v2.0.87)(Tischler & Leonard, 2014); non-duplicate, properly paired reads were kept by samtools (parameter “-q 1 -F 1804” version 1.2)(H. Li et al., 2009). To adjust Tn5 shift, reads were offset by +4 bp for the sense strand and −5 bp for the antisense strand. We used fragment size to separate reads into nucleosome-free, mononucleosome, dinucleosome, and trinucleosome groups as described (Buenrostro et al., 2013) and generated bw files by using the center 80 bp of fragments and scale to 20 million nucleosome-free reads. Two replicates were merged to enhance peak calling by MACS2 (version 2.1.1.20160309 default parameters with “--extsize 200 -- nomodel “)(Zhang et al., 2008). All of the important nucleosome-free regions were considered to have been called if a sample had more than 15 million nucleosome-free reads after merging. To ensure reproducibility, we first merged peaks from different cell types to create a set of reference chromatin-accessible regions. We then used bedtools (v2.24.0)(Quinlan & Hall, 2010) to count nucleosome-free reads from each sample to overlay with the reference regions. To identify differentially accessible regions, we first normalized raw nucleosome-free reads counts by using trimmed mean of M-values (TMM) and then applied empirical Bayes statistics testing after linear fitting with voom package (R 3.23, edgeR 3.12.1, limma 3.26.9)(Law, Chen, Shi, & Smyth, 2014). A false discovery rate (FDR)–adjusted *p*-value <0.05 and fold change > 2 were used as cutoff values for differentially accessible regions.

### ChIP qPCR and sequencing

CD4 Tn cells from *Foxp3*^*gfp*^ male mice were induced to become iT_reg_ or Th0 cells in the presence or absence of 0.25 mM ascorbic acid-2-phosphate. The nT_reg_ cells from *Foxp3*^*gfp*^ male mice were FACS sorted and expanded *in vitro* for 5 days by using Dynabeads Mouse T-Activator CD3/CD28 (Invitrogen) in the presence of 500 U/mL recombinant IL-2. To perform Stat5 ChIP, iT_reg_ or nT_reg_ cells were starved for 3 h in complete RPMI1640 without cytokines. Cells were then restimulated with 500 U/mL IL-2 for 30 min. A two-step protocol was used to fix the cells as we previously described (Feng et al., 2014). In brief, cells were resuspended at 5×10^6^ cells/mL in 1 mM MgCl_2_/PBS and treated with 2 mM DSG cross-linker (Thermo Fisher Scientific) at room temperature for 30 min on a rotator. Cells were then washed twice with PBS and fixed with 1% formaldehyde at room temperature for 5 min. Fixation was quenched by adding 125 mM glycine. Cells were washed once with ice-cold PBS, aliquoted, and either frozen at − 80°C or immediately processed by downstream reactions. Chromatin sonication was performed by using the truChIP Chromatin Shearing Kit (Covaris) with Focused-Ultrasonicator M220 (Corvaris) following the manufacturer’s instructions. Chromatin was sheared to 400-800 bp for Stat5 and Foxp3 ChIP; 10% of samples were aliquoted as input control. In each ChIP reaction, 5 µL of rabbit anti-Stat5 (CST), 10 µL of rabbit anti-Foxp3 (Abcam), or control rabbit IgG (CST) were added to the lysis/binding buffer (50 mM HEPES pH 8.0, 1 mM EDTA, 1% Triton X-100, 0.1% sodium deoxycholate, 0.1% SDS, 140 mM NaCl, 1 mM PMSF, and protease inhibitor cocktail) to precipitate the chromatin. After an overnight incubation, protein A and protein G magnetic beads (Invitrogen) were added to capture the antibody-chromatin complexes. Next, beads were washed according to our published protocols (Feng et al., 2014). To release the DNA, ChIP samples were treated with proteinase K, followed by phenol:chloroform:isoamyl alcohol extraction and 2-propanol precipitation in the presence of GlycoBlue Coprecipitant (Invitrogen). DNA pellets were dissolved in 1×TE buffer (10 mM Tris HCl pH 8.0, 1 mM EDTA) for qPCR or deep sequencing.

To quantify the precipitated DNA, qPCR was performed with locus- or region-specific primers, PowerUp SYBR Green Master Mix (Applied Biosystems), and the CFX384 Real Time PCR System (Bio-Rad). Relative enrichment of the targets was calculated by normalizing the signals of the precipitated DNA to those of the input samples. To perform ChIP-Seq, libraries were prepared with precipitated DNA by using the KAPA Hyper Prep Kit (Kapa Biosystems). The library DNA was enriched by size selection with AMPure XP beads and quantified by using the NEBNext Library Quant Kit. Indexed samples were pooled together for 100 cycles of paired end sequencing on a HiSeq 4000 or HiSeq 2500 system (Illumina). More than 40 million reads per sample were sequenced for downstream analysis.

Single-end reads of 50 bp were mapped to mouse genome mm9 (MGSCv37 from Sanger) by BWA (version 0.7.12-r1039, default parameter)(H. Li & Durbin, 2009). Duplicate reads were marked with biobambam2 (version v2.0.87)(Tischler & Leonard, 2014); non-duplicate reads were kept by samtools (parameter “-q 1 -F 1024” version 1.2) (H. Li et al., 2009). We followed ENCODE guidelines to assess the quality of our data as previously described(Yang et al., 2019). Next, we extended reads to fragment size (detected by SPP v1.1)(Kharchenko, Tolstorukov, & Park, 2008) and generated bw tracks (normalized to 10 million uniquely mapped reads) for visualization. We used MACS2 (version 2.1.1.20160309, parameters “--nomodel --extsize fragment size”) to call peaks. To assure reproducibility, we finalized the peaks for each group if they were called with a stringent cutoff value (FDR-adjusted *p*-value < 0.05 in MACS2) in one sample and at least called with a lower cutoff value (FDR-corrected *p*-value < 0.5 in MACS2) in the other. Peaks were further merged between groups. Reads were extended to fragment size for each sample via bedtools (v2.24.0)(Quinlan & Hall, 2010). We used correlation plots to assess reproducibility among replicates. After TMM normalization, we used empirical Bayes statistics testing after linear fitting from voom package(R 3.23, edgeR 3.12.1, limma 3.26.9)(Law et al., 2014) to identify differential binding sites. FDR-adjusted *p*-value <0.05 and fold change > 2 were used as cutoff values.

### Quantification of nascent RNA

Cells were labeled with 0.5 mM ethylene uridine for 45 min (2.5 × 10^6^ cells/mL culture medium) before being lysed with TRIzol according to the manufacturer’s protocol (Thermo Fisher Scientific). Total RNA was isolated via chloroform separation and 2-propanol precipitation; 1 µg RNA was then used to perform the click reaction followed by a purification with Dynabeads MyOne Streptavidin T1 beads according to the manufacturer’s protocol (Click-iT Nascent RNA Capture Kit, Thermo Fisher Scientific). Enriched nascent RNA was immediately used as a template to generate cDNA with the SuperScript VILO cDNA Synthesis Kit (Thermo Fisher Scientific). Quantification of Foxp3 transcripts was conducted with specific primers on the CFX384 Real Time PCR System (Bio-Rad).

### Mixed bone marrow chimeras

Mixed bone marrow chimeric mice were generated as we previously described (Feng et al., 2015). Briefly, recipient mice (CD45.1^+^CD45.2^+^) were lethally irradiated (9.5 Gy) 24 h before intravenous injection of 5 × 10^6^ to 10 × 10^6^ bone marrow cells from CD45.1 WT and CD45.2 *Foxp3*^*LSL*^ mixed at a 1:1 ratio. After bone marrow transfer, the recipient mice were administered neomycin (2 mg/mL) in drinking water for 3 weeks and euthanized for CD4 T cell preparations 8-10 weeks later.

### Cut&Run experiments

Viable iT_reg_ cells were sorted by *Foxp3*^*gfp*^ reporter expression after induction. Cut&Run was performed as described (Skene & Henikoff, 2017). Briefly, 1×10^6^ iT_reg_ cells were first attached to Concanavalin A (ConA)-coated magnetic beads, permeabilized with digitonin-wash buffer (20 mM HEPES, pH7.5, 150 mM NaCl, 0.5 mM spermidine, 0.01% Digitonin, and protease inhibitors), and then incubated with antibody (1:100 H3K27ac) for 2 h at 4°C on a rotator. The beads-cell mixture was washed thrice with digitonin-wash buffer, resuspended in 200 µL of protein A-MNase, and incubated for 1 h at 4°C on a rotator. After 3 rounds of washing, beads were resuspended in 150 µL of digitonin-wash buffer and chilled in an ice-water bath for 5 min; 3 μL of 100 mM CaCl_2_ was added into the tubes with gentle vortexing, and the beads were incubated in an ice-water bath for 30 min. Next, 150 µL of 2×STOP buffer (170 mM NaCl, 20 mM EDTA, 20 mM EGTA, 0.05% Digitonin, 20 mg/mL GlycoBlue, 25 mg/mL RNase A) was added and incubated at 37°C for 30 min. After clarification on a magnet stand, supernatant was transferred to a fresh tube for phenol:chloroform extraction and ethanol precipitation. A library was prepared by using KAPA Hyper Prep Kit, and 5ng DNA was loaded for deep sequencing.

After adapter trimming by cutadapt (version 1.9, paired-end mode, default parameter with “ -m 25 -O 6 “)(Martin, 2011), 50-bp paired-end reads were mapped to mouse genome mm9 (MGSCv37 from Sanger) by using BWA (version 0.7.12-r1039, default parameter)(H. Li & Durbin, 2009). Duplicate reads were marked with biobambam2 (version v2.0.87)(Tischler & Leonard, 2014); non-duplicate reads were kept by samtools (parameter “-q 1 -F 1804” version 1.2)(H. Li et al., 2009). At least 5 million reads per sample were used for further analysis, as suggested in published research (Skene, Henikoff, & Henikoff, 2018). We generated bw files using the center 80 bp of fragments smaller than 2000 bp and normalizing the reads counts to 10 million fragments.

### RNA sequencing

T_reg_ cells induced from male WT *Foxp3*^*gfp*^ CD4 Tn cells in the presence or absence of 0.25 mM ascorbic acid-2-phosphate were sorted on day 4 during culture. Cells were lysed with TRIzol; RNA was extracted, and sequencing libraries were constructed by using SMARTer cDNA synthesis kits (Takara). Libraries were sequenced by using the HiSeq 2500 platform (Illumina) according to a standard paired-end protocol. Reads processing, mapping, and differential gene expression analyses were performed as we previously described^25^.

### Data resources

The sequencing data have been deposited at https://www.ncbi.nlm.nih.gov/geo/ with access number GSE146250.

### Software resources

The R codes used for the analyses are available upon request.

